# A rationally designed mimotope library for profiling of the human IgM repertoire

**DOI:** 10.1101/308973

**Authors:** Anastas Pashov, Velizar Shivarov, Maya Hadzhieva, Victor Kostov, Dilyan Ferdinandov, Karen-Marie Heinz, Shina Pashova, Milena Todorova, Tchavdar Vassilev, Thomas Kieber-Emmons, Leonardo A. Meza-Zepeda, Eivind Hovig

**Author notes:** Corresponding author. **Mailing Address:** Institute of Microbiology, BAS, Acad. G Bonchev St, block 26 Sofia 1113, Bulgaria **E-mail:** (AP).

## Abstract

Specific antibody reactivities are routinely used as biomarkers but the use of antibody repertoire profiles is still awaiting recognition. Here we suggest to expedite the adoption of this class of system level biomarkers by rationally designing a peptide array as an efficient probe for an appropriately chosen repertoire compartment. Most IgM antibodies are characterized by few somatic mutations, polyspecificity and physiological autoreactivity with housekeeping function. Previously, probing this repertoire with a set of immunodominant self-proteins provided only coarse information on repertoire profiles. In contrast, here we describe the rational selection of a peptide mimotope set, appropriately sized as a potential diagnostic, that also represents optimally the diversity of the human public IgM reactivities. A 7-mer random peptide phage display library was panned on pooled human IgM. Next generation sequencing of the selected phage yielded a non-exhaustive set of 224087 mimotopes which clustered in 790 sequence clusters. A set of 594 mimotopes, representative of the most significant clusters, was used to demonstrate that this approach samples symmetrically the space of IgM reactivities. When probed with diverse patients’ sera in an oriented peptide array, this set produced a higher and more dynamic signal as compared to 1) random peptides, 2) random peptides purged of mimotope-like sequences and 3) mimotopes from a small subset of clusters. In this respect, the representative library is an optimized probe of the human IgM diversity. Proof of principle predictors for randomly selected diagnoses based on the optimized library demonstrated that it contains more than 10^70^ different profiles with the capacity to correlate with diverse pathologies. Thus, an optimized small library of IgM mimotopes is found to address very efficiently the dynamic diversity of the human IgM repertoire providing informationally dense and structurally interpretable IgM reactivity profiles.

**Author Summary:** The presence in the blood of antibodies specific for a particular infectious agent is used routinely as a diagnostic tool. The overall profile of available antibody reactivities (or their repertoire) in an individual has been studied much less. As an omics approach to immunity it can be a rich source of information about the system beyond just the individual history of antigenic exposure. Using a subset of antibodies – IgM, which are involved also in housekeeping functions like removing dead cells, and bacteriophage based techniques for selection of specific peptides, we managed to define a non-exhaustive set of 224087 peptides recognized by IgM antibodies present in most individuals. They were found to group naturally in 790 structural groups. Limiting these to the most outstanding 594 groups, we used one representative from each group to assemble a reasonably small set of peptides that extracts the maximum information from the antibody repertoire at a minimum cost per test. We demonstrate, that this representative peptide library is a better probe of the human IgM diversity than comparably sized libraries constructed on other principles. The optimized library contains more than 10^70^ different potentially profiles useful for the diagnosis, prognosis or monitoring of inflammatory and infectious conditions, tumors, neurodegenerative diseases, etc.

## Introduction

The repertoire of human IgM contains a considerable proportion of moderately autoreactive antibodies characterized by low intrinsic affinity/ low specificity, functioning as a first line of defense [1], as scavengers of senescent cells and debris [2–6], and even in tumor surveillance [7]. It is becoming increasingly clear that the human antibody repertoire has an organization similar to that of its murine counterpart [8–12]. About one fourth of the murine splenic B lymphocytes that respond to lipopolysaccharide have B cell receptors which are moderately autoreactive. Affected very little by somatic mutations and follicular evolution, the physiological self-reactivities largely overlap with germline-encoded polyspecific antibodies [13–15]. Eighty percent of murine serum IgM falls in this category and is referred to as natural antibodies (nAbs) [16,17]. Apart from the polyspecific splenic B cells, the source of nAbs in mice seems to be mostly a population of B1-related IgM^+^ plasma cells residing in a unique IL-5 dependent bone marrow niche [18].

The IgM antibody repertoire is an insufficiently explored source of biomarker profiles. IgM antibodies appear early in the course of an infection. However, they fall relatively fast, even after restimulation, providing a dynamic signal. By interacting with structures of self and carrying housekeeping tasks, this part of the antibody repertoire reacts swiftly to and reflects changes in the internal environment. Consequently, IgM antibodies have gained interest as biomarkers of physiological or pathological processes [19–23], but remain underused as immunodiagnostics, although their interactions with sets of antigens have been studied in a range of platforms [19,22–25].

The study of the IgM repertoire might be expected to give information about interactions that occur mostly in the blood and the tissues with fenestrated vessels since, unlike IgG, IgM cannot easily cross the normal vascular wall. Yet, IgM tissue deposits are a common finding in diverse inflammatory conditions [26–28] and especially in the disorganized vasculature of the tumors, where they are a key element of the innate immune surveillance mechanism [7,29,30]. Changes in the IgM repertoire further reflect B cell function affected by antigenic, danger and inflammatory signals, but also by anatomical changes leading to vascular permeability or disruption. Thus, IgM repertoire monitoring has the potential to provide clinically relevant information about most of the pathologies involving inflammation and vascular remodeling, as well as all types of cancer.

Our working hypothesis is that an essential part of the human IgM repertoire involved in homeostasis can be probed by a set of mimotopes, the size of which can be tailored to the diagnostic goals by optimization. The existing approaches for immunosignature [31,32] or immunomic [33] analysis of the immunoglobulin repertoires focus mostly on IgG and have used arrays of either 10^2^ proteins or 10^4^-10^5^ random peptides. The IgM repertoire has been previously probed by protein arrays [34] containing a biologically determined representative set of autoantigens which is a structurally coarse approach. We set out to explore the feasibility of a method that, similar to the self-protein “homunculus” arrays, targets a small set of rationally selected probes, but also preserves the structural interpretability of peptides in a format applicable for routine diagnostics.

## Results

### Selection of 7-mer mimotopes

We chose to pan a commercially available 7-mer random peptide phage display library of diversity 10^9^. Thus, the size of the mimotopes would be in the range of the shorter linear B cell epitopes in the IEDB database (http://www.iedb.org/). At the same time, the almost complete diversity of sequences of that length could be interrogated. As a repertoire template we used an experimental preparation of human immunoglobulins for intravenous use enriched in IgM representing a pool of the repertoire of approx. 10 000 healthy donors. The phage eluted from the IgM repertoire were adsorbed on a monoclonal IgM to filter out phage binding to the constant regions, and thereby focus only on the mimotopes (Fig. 1). The peptide inserts were amplified and deep sequenced using the approach described by Matochko et al. (2012) [35]. Two separate experiments starting with 20% of the original phage library were performed (experiments A and B), while in a third one (C), a preamplified 20% sample of the original phage library was used. The yield was 688 860 (experiment A), 518 533 (experiment B) and 131 475 (experiment C) unique reads. Based on the distribution of the reads by copy number in the selections from the native and preamplified library two thresholds were determined – 2 and 11 copies, and the reads within these limits were considered further (see Suppl. Methods).

**Figure 1.**
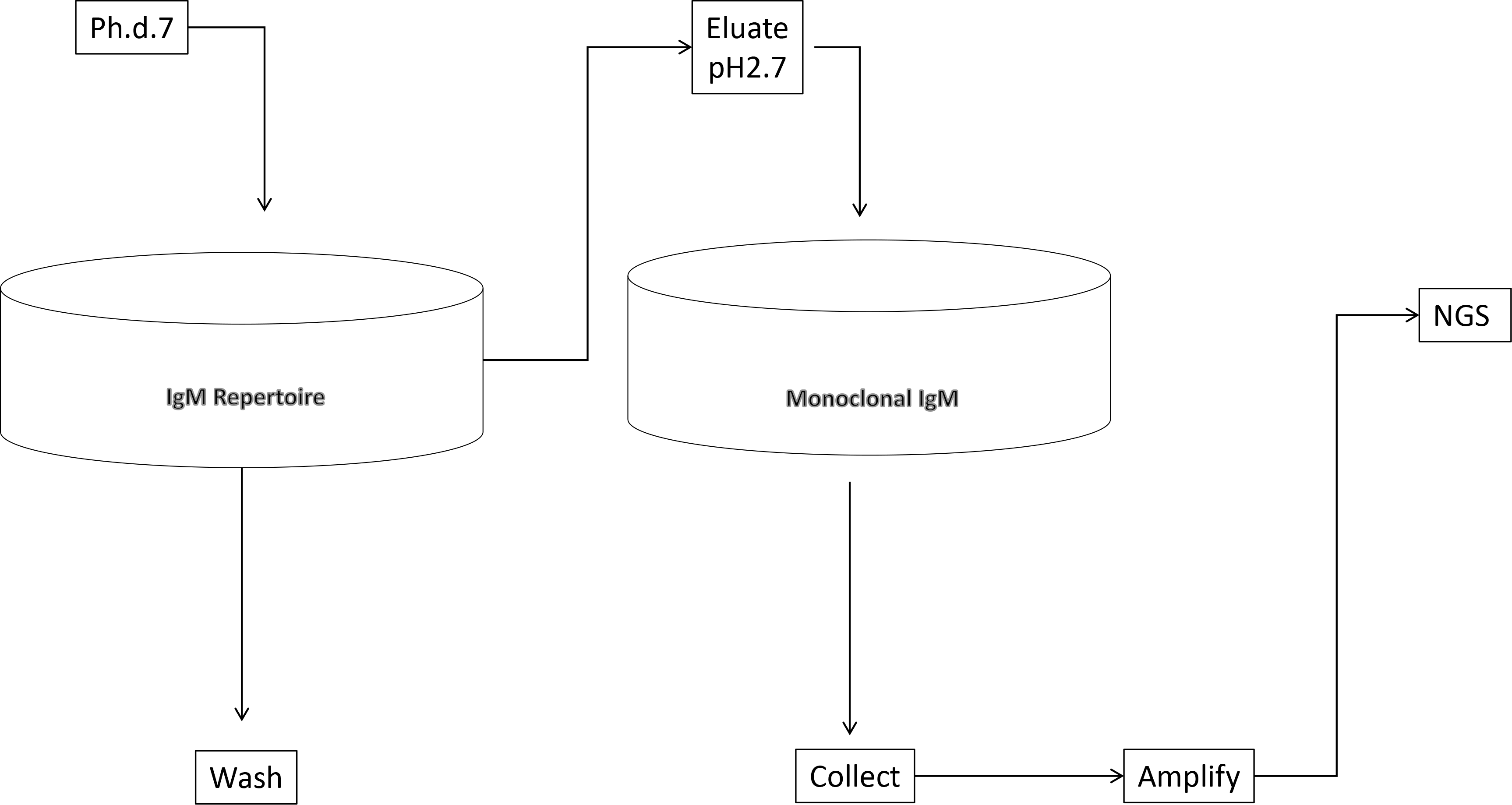
Schematic representation of the deep panning experiment.

### Sequence properties of the mimotope clusters

The overall amino acids residues frequencies (AAF) in the mimotopes selected from the phage library showed a skewing in favor of G,W,A,R,T,H,M,P,Q and against C,F,N,Y,I,L and S (Fig. 2A) when compared to the average overall amino-acid frequencies of the Ph.D.-7 library. When studied by position, the distribution of AAF visualized by the respective sequence logos showed a highly skewed distribution, diverging from the overall background frequencies, only for the N-terminus (Fig. 2B). The actual frequencies by position are shown in Fig. 2C. The residues of W, D and E appear in similar frequencies but due to the much lower abundance of W, generally and in the phage library, in Fig. 2A it comes up as selected and D and E – as slightly disfavored. Somewhat surprisingly, the N terminal frequencies skewing and the preference for A, P and T proved to be properties of the library when comparing the AAF by position of a non-selected but amplified library (based on the data from Matochko et al., Fig. 2D). The evidence of selection by IgM stood out in the distribution by position only after using the PWM of the non-selected amplified library as background frequencies to described the actual enrichment in our mimotope library (Fig. 2E). It showed higher divergence from the background distribution of the frequencies in the middle of the sequence. Overrepresentation of proline in positions 2-7 appears to be a property of the amplified library (background frequencies plus collapse of diversity) while the IgM binding selected for negatively charged residues, glycine and tryptophan.

**Figure 2.**
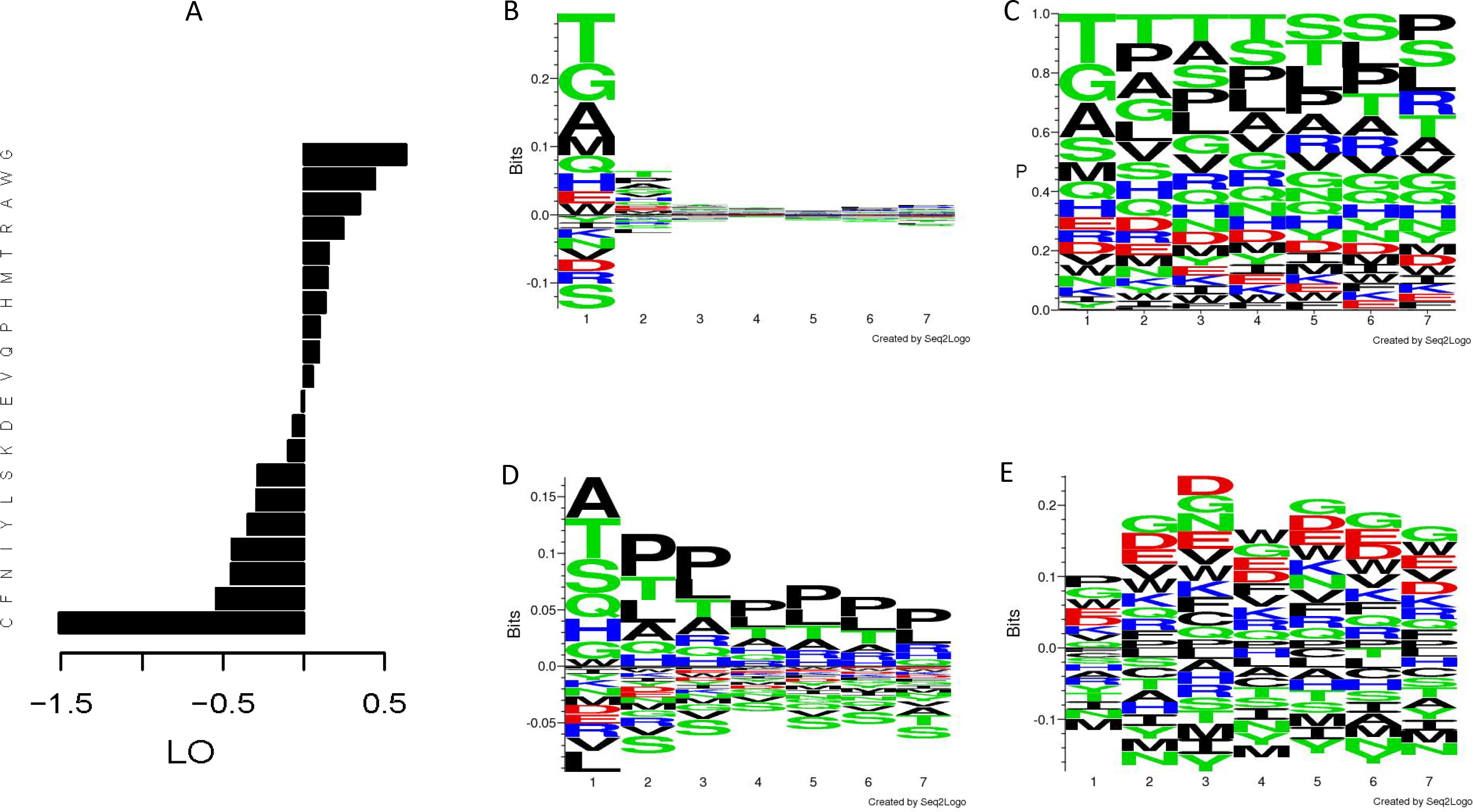
Distribution of the amino acid residues in the mimotope library. (A) Log odds (LO) relative to background frequencies; (B) sequence logo of the LO by position relative to the overall background frequencies; (C) sequence logo of the frequencies by position; (D) sequence logo of the LO by position in an amplified Ph.d.-7 phage library without ligand selection relative to the overall background frequencies (based on W. L. Matochko, R. Derda. “Error analysis of deep sequencing of phage libraries: peptides censored in sequencing”. Comput Math Methods Med, 2013, 491612. 2013., http://www.chem.ualberta.ca/~derda/parasitepaper/rawfiles/PhD7-GAC-30FuR.txt); (E) sequence logo of the LO by position relative to the frequencies by position in the amplified, unselected library shown in (D). The skewing of the distribution in the free N-terminus appears to be property of the library while the selection by the IgM repertoire leads to slight skewing in the middle of the sequence towards negatively charged residues, glycine and tryptophan.

To gain insight into the mimotope sequence space the set of 224 087 selected mimotope sequences was subjected to clustering using the GibbsCluster-2.0 method [36] originally applied for inferring the specificity of multiple peptide ligands tested on multiple MHC receptors. The number of clusters was optimized in the range of 100 to 2500 clusters using the Kullback-Leibler divergence criterion (an information theory based measure of similarity between two distributions, in this case – two sequence profiles) comparing the sequences to the background model of random sequences [36]. This criterion indicated optimal clustering in 790 clusters (Fig. 3). Position weighted matrices (PWM) were calculated from each cluster (Supplement file 2).

**Figure 3.**
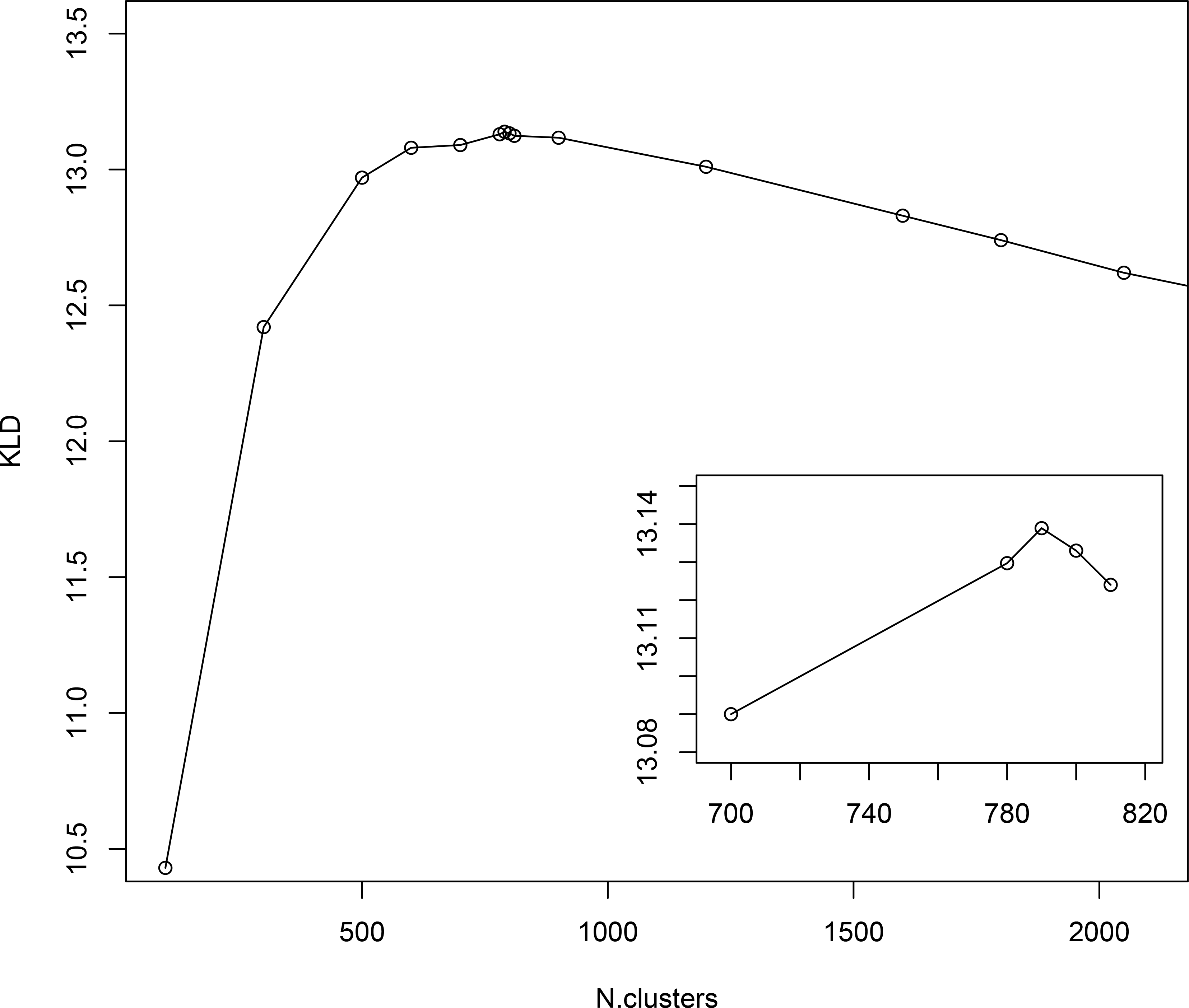
Results from GibbsCluster of the mimotopes. Different predefined number of clusters were screened for the quality of clustering measured by Kullback-Leibler Divergence – KLD. The inset shows amplified scale around the peak KLD values.

### Generation of libraries of 7-mers targeting different aspects of the IgM igome

The mimotope library of more than 200 000 sequences is a rich source of potential mimotope candidates for vaccine or diagnostics. The relevance of this mimotope library to the complete IgM repertoire and the scope of its diversity could be probed comparing several different peptide libraries with different properties. An alternative library was constructed to check the completeness of the selected mimotope set (what part of the igome it represents) and the relevance of the clustering found. To this end, 2.3×10^6^ random 7-mer sequences were scored for their similarity to each cluster profile and ranked. The random sequences that were the least related to any of the clusters in the selected library were used as a negative control (library pepnegrnd – see Suppl. Methods).

As a probe of the IgM repertoire for routine diagnostic use, an array of 10^5^ peptides is of an impractical size. A way to construct an optimal smaller mimotope library would be to include a representative sequence of each of the naturally existing 790 clusters as they would sample evenly (symmetrically) the mimotope sequence space as ensured by the GibbsCluster algorithm. The clusters were found to vary with respect to the probability of random occurrence so subsequently they were ranked by significance using this probability (Suppl. Methods). The top 594 clusters were considered further and only the mimotope with the top score from each cluster was kept as a mimotope prototype for the profile. This library was labeled peppos.

Other libraries of peptides generated for further comparison were: 1) uncertainly clustered sequences as reflected in their Kulback-Leibler Divergence scores as shown by the GibbsCluster algorithm (pepneg and pepneglo); 2) 2 groups of 5 highest scoring clusters – lower diversity libraries (pep5 and pepother5); random 7-mer sequences predicted to belong to any of the 5 highest scoring clusters based on profile scores (pep5pred) and 4) random 7-mer sequences (peprnd) (see Table 1 for description of all libraries). The number of sequences per library was constrained by the size of the chip.

**Table 1.**
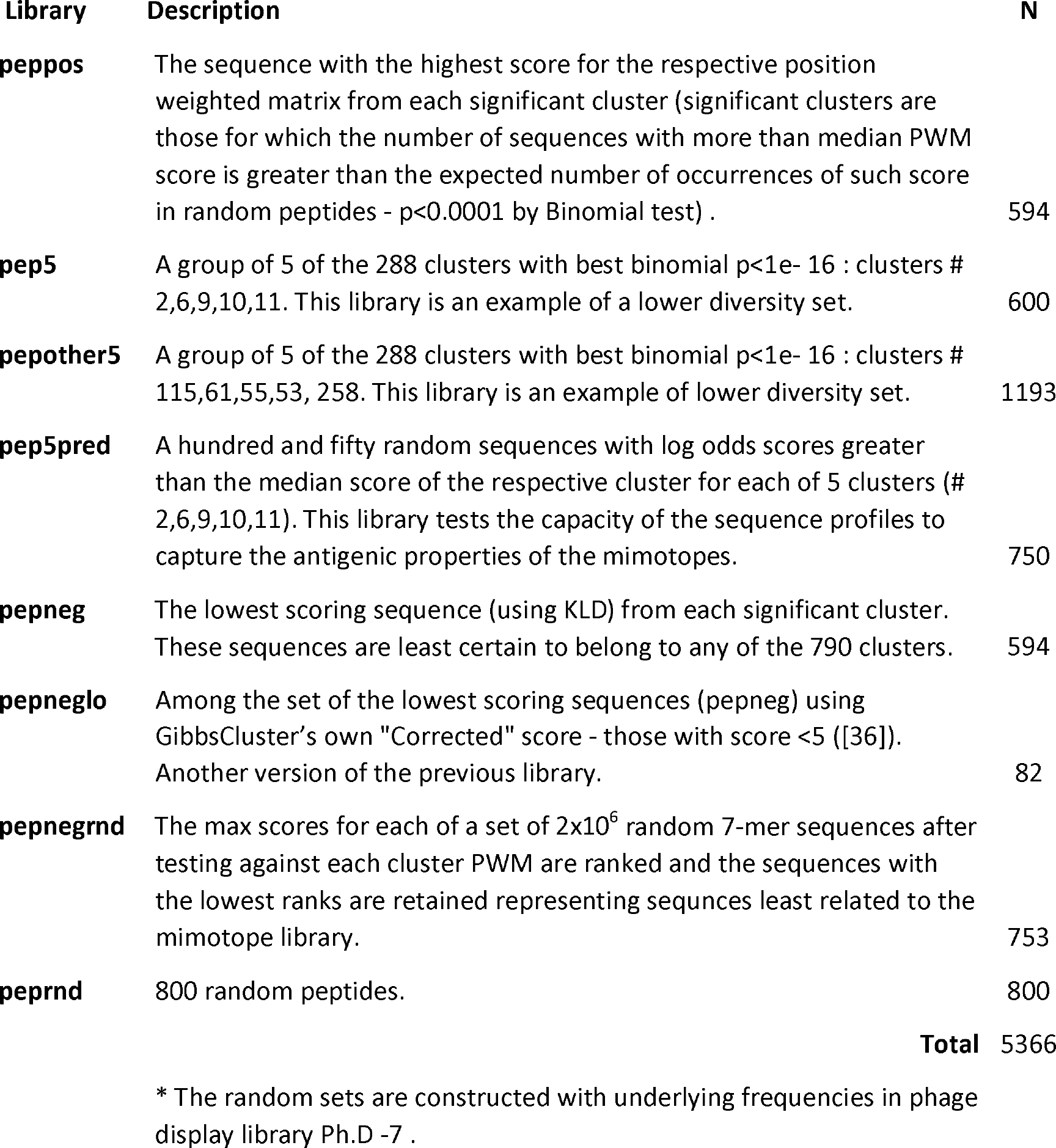
Libraries of 7-mer peptides studied

### Comparison between libraries

IgM reactivity in sera from patients with glioblastoma multiforme (GBM), brain metastases of breast cancers (MB) as well as non-tumor bearing neurosurgery patients (C) was analyzed using the sets of peptides described in Table 1. The peptide libraries were synthesized in an oriented (C-terminus attached) planar microarray format. In the first round of experiments, the 8 different libraries defined were compared based on the IgM reactivity in the sera from 10 patients (Suppl. Fig. 4 and 5). The data on the mean serum IgM reactivity of the peptides, grouped by library, with the different sera was used to compare the libraries for their overall reactivity using linear models (Fig. 4A). The proposed optimized small library (peppos) had significantly higher (p<0.001) average reactivity than pepneg, peprnd or pepnegrnd. Interestingly, the library theoretically purged of relevant reactivities (pepnegrnd) had indeed the lowest reactivity, significantly lower than both the weakly clustering peptides (pepneg) and the random sequences (peprnd) (Suppl. Table 1).

Next, the capacity of the different libraries to sample symmetrically the space of mimotope reactivities was tested. To this end, the total correlation of the IgM reactivity profiles of the peptides in each library mapped on the ten different sera (Fig. 4B) were compared. The total correlation is a KLD based multidimensional generalization of mutual information. High total correlation would signify redundancy in the library with many peptides sharing similar reactivity profiles. The library peppos had the lowest total correlation, while pepnegrnd, and especially pep5pred had the top total correlation, indicating redundancy of the information their reactivity carries about the patients, while the pools of 5 clusters – pep5 and pepother5 – had relatively low correlation. All differences, except between the top two libraries were significant (Suppl., Table 2).

Another way to test the symmetry of the representation of the mimotope reactivity space by the different libraries is to compare the mean nearest neighbor distance (MNND) of the scaled and centered data of mimotope staining intensity mapped again to the 10 patients’ sera IgM reactivity. Peptides which have similar reactivity profiles with different sera would map to points in the reactivity space that are close to each other. This clustering in some regions of the space would lead to a lower MNND. The library peppos ranked second only to pepneglo (Fig. 4D) by this parameter and had a significantly higher MNND than all the other libraries (Suppl. Table 4.).

The correlation of the serum profiles based on the different mimotopes (transposing the matrices of the previous tests) can also be viewed as a criterion for the capacity of the libraries to extract information from the IgM repertoire. Due to the extreme multidimensionality, the mean correlation between patient profile pairs was used to compare the libraries after z transformation of the correlation coefficient to allow comparison by linear models (Fig. 4C). Again, the peppos library exhibited the lowest mean correlation - significantly lower compared to the correlation between the reactivities with the other libraries except for pepnegrnd and pepneglo (Suppl. Table 3.)

**Figure 4.**
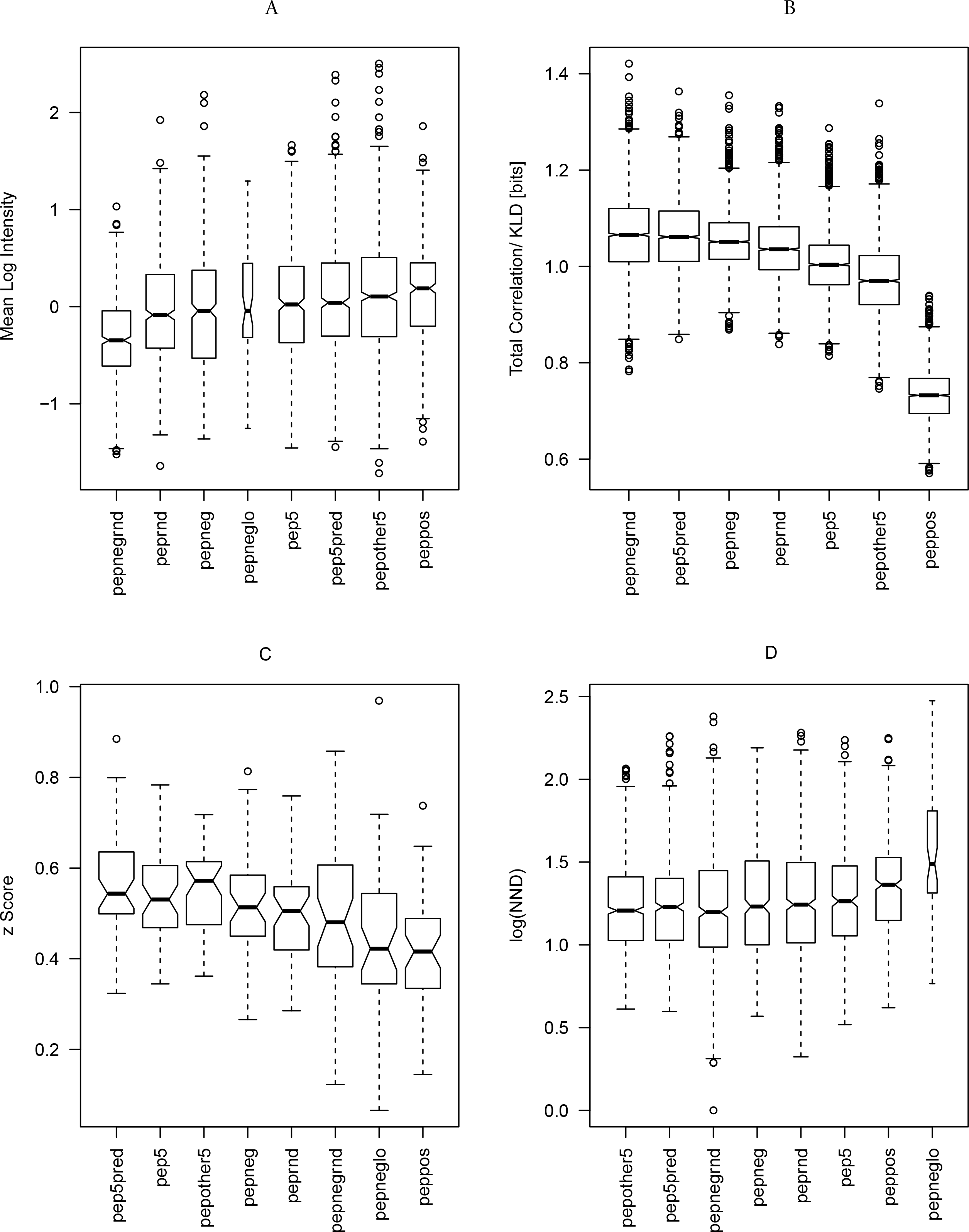
Statistics testing the libraries’ capacity to probe the mimotope reactivity space. A) Mean reactivity of each peptide across patients grouped by library. The optimized library peppos has the highest reactivity. For library content see Table 1. B) Total correlation of the peptide profiles grouped by library across 10 patients. The optimized library peppos provides the least redundant information. C) Mean correlation of patient profiles across the peptides in each library compared after z-transformation. The optimized library peppos provides the most diverse characteristics of the patients, which indicates a high potential for discrimination of different states but increases the requirements for the size of the teaching sets to extract models of good generalization. D) Mean nearest neighbor distance of the peptide profiles across 10 patients in each library. Again, the optimized library peppos appears to sample the mimotope reactivity space evenly. The width of the bars is proportional to the size of the sample.

Finally, all four criteria were summarized using a rank product test, which proved that reactivity with peppos stands out from all the other tested libraries as the best among them for probing the IgM repertoire (Table 2).

**Table 2.**
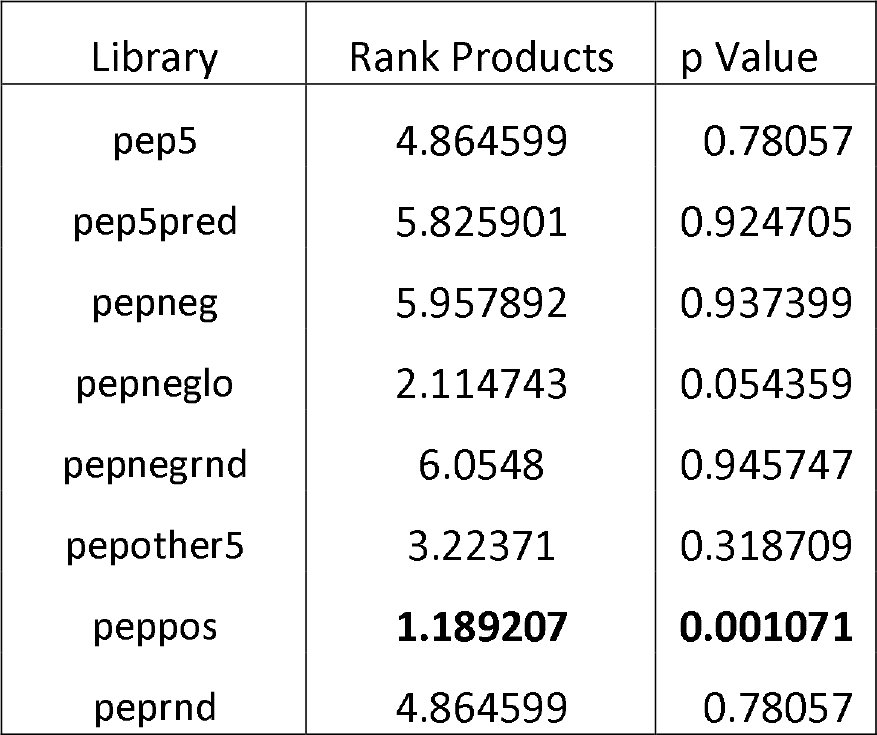
Rank product test of four criteria for optimal mimotope library:

### Visualization of the Mimotope Space

T distributed stochastic neighbor embedding (t-sne) was used to visualize the structure of the mimotope sequence space as represented by the general mimotope library produced by deep panning. To represent the sequences as vectors of real numbers, each amino acid residue was represented by 5 scores based on the z1-z5 scales published by of Sandberg et al. (1998) (see Suppl. Methods for details). Thus, each 7-mer sequence was parametrized as a 35-dimensional vector. These vectors were then represented in two dimensions by t-sne transformation. The map of the mimotope library, thus generated, resembled that of an equal number of random 7-mer sequences constructed using the residue background frequencies of the phage display library (Suppl. Fig. 7). Next the representation of some of the clusters of mimotopes described above were mapped in this new mapping. Although most of the five most significant among 790 sequence clusters (the pep5 library, Suppl. Figure 7) mapped to rather scattered clusters, the mapping of the optimized library (peppos) still covered symmetrically the mimotope sequence space (Fig. 5). Both the clustering and the mapping do not give unique solutions and fail to capture the full information in the general mimotope data set. Yet, the symmetry of the rationally small library designed on the basis of the clustering is preserved in the t-sne mapping indicating that it is an actual property of the small library peppos.

Mapping together mimotopes and random peptides helped estimate the proportion of the 7-mer sequence space which is under sampled by the mimotope library (Fig. 6, see Suppl. Methods for details). To partition the sequences in logical groups, k-means clustering on the t-sne map was performed. The proportion of mimotopes and random sequences in each cluster were calculated next. While all clusters defined by the k-means clustering contained mimotopes, the proportion of mimotope varied producing 2 types of clusters – some with equal representation of random sequences and mimotopes and some with predominance of random sequences. The random peptide sequences with minimal similarity to the IgM igome (library pepnegrnd) mapped also to the areas of the low density of mimotopes. Approximately 42 % of the random sequences, 14% of the mimotopes and 85% of the pepnegrnd library were in the underrepresented areas (Chi square, p<0.0001). The areas of the sequence space underrepresented in the IgM mimotopes had very similar sequence profiles to the normally represented areas, except for less abundant charged residues (Suppl. Data file “t-sne cluster profiles”).

**Figure 5.**
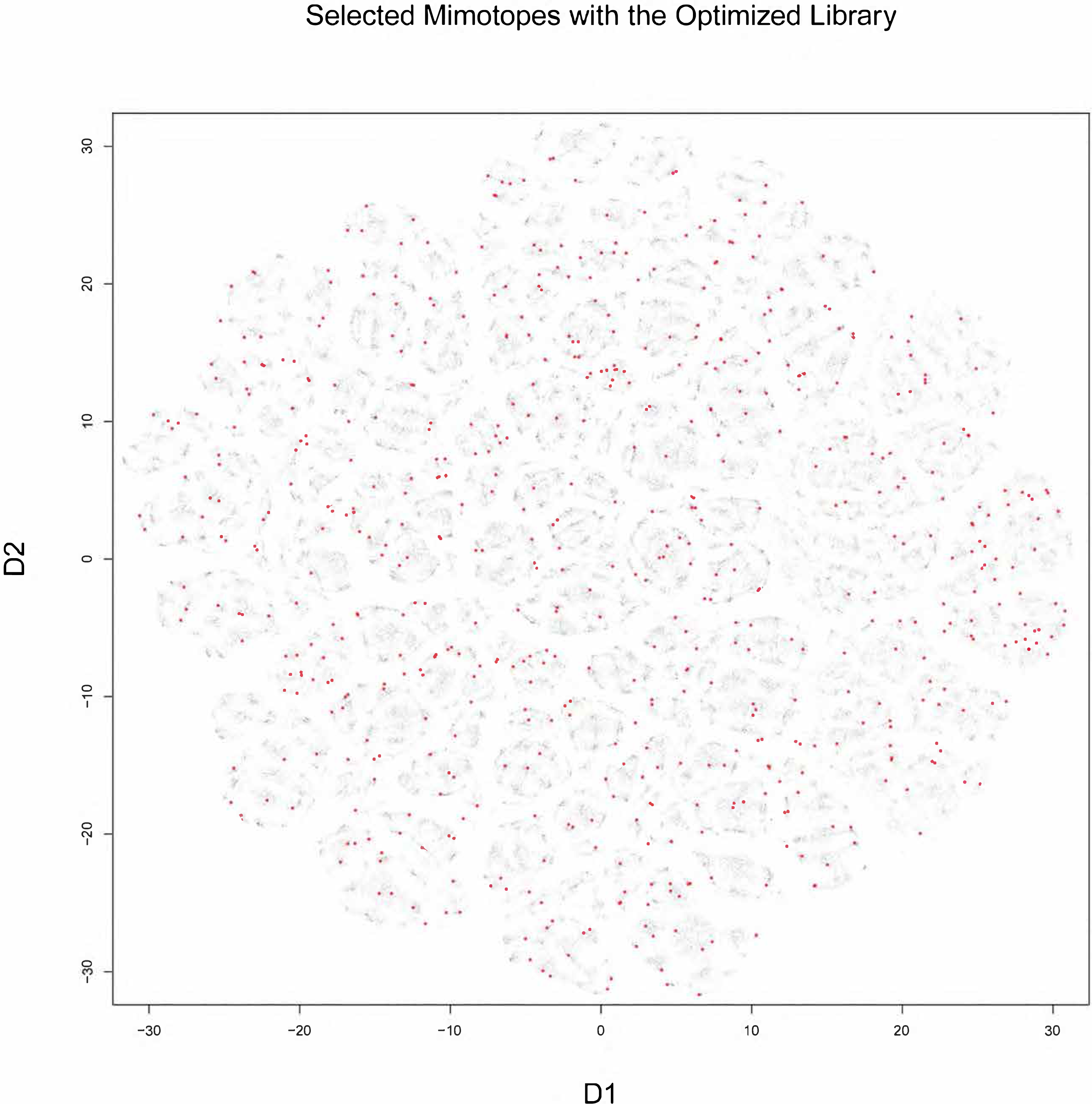
Visualization of the 7-mer mimotope sequence space with the optimized library peppos marked in red (see fig.8 for details). Although, individual GibbsCluster defined clusters do not coincide with those shown by t-sne, the mapping of the optimized library apparently probes quite uniformly the mimotope sequence space.

**Figure 6.**
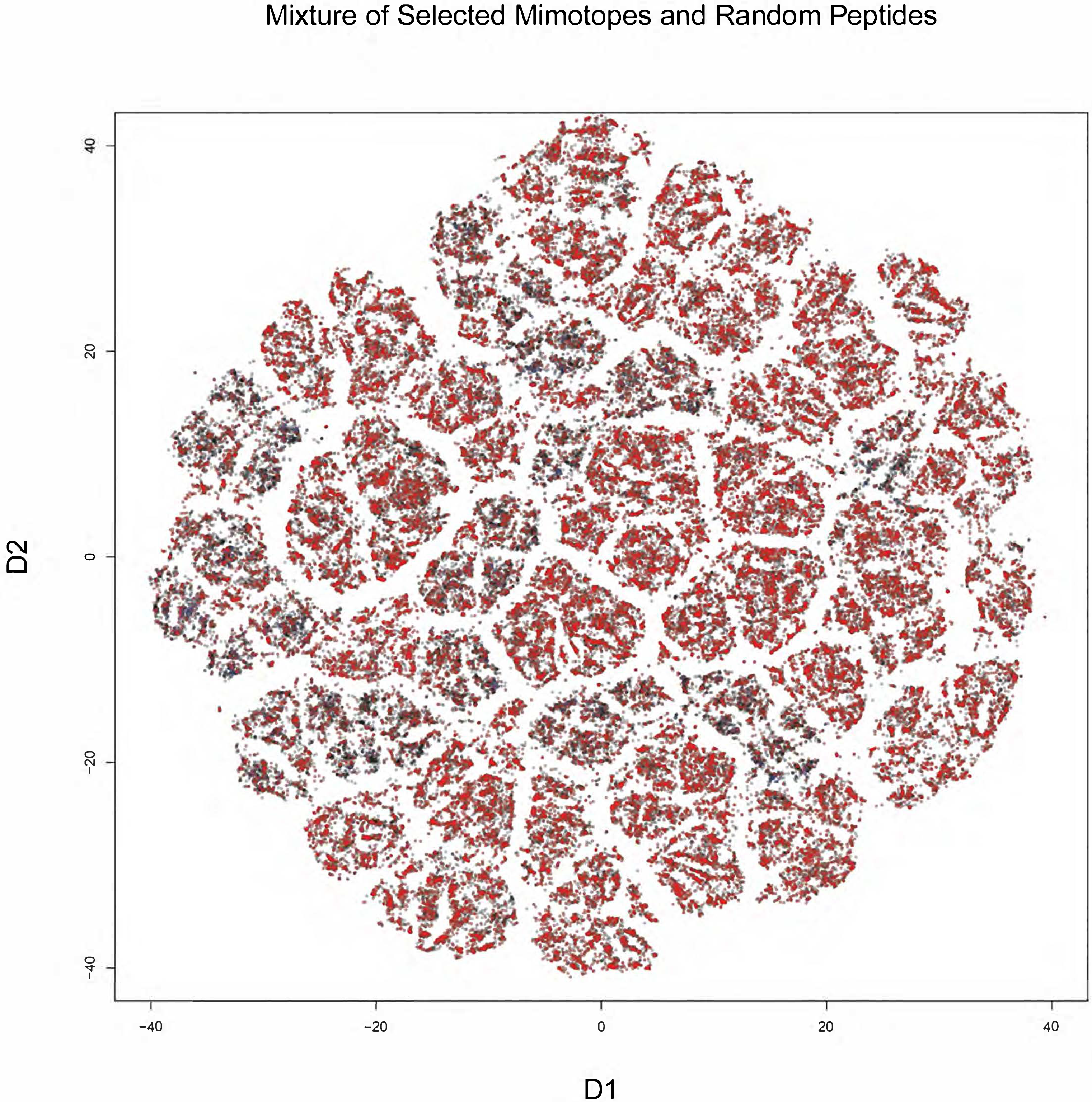
Visualization of the 7-mer peptide mimotope sequence space representing a mixture of random sample of 50 000 phage display selected mimotopes (red) and 50 000 random sequences (gray) plus the pepnegrnd library (blue). A part of the sequence space is represented by mimotopes at a lower density and the sequences unrelated to the defined 790 mimotope clusters map mostly to this area (blue points). A high definition version of this figure is included in the supplemental information.

### Diagnostic potential of a rationally designed restricted mimotope library

A suitably sized universal mimotope library sampling optimally the public IgM reactivities would have multiple applications both in the theoretical research of antibody repertoires, as well as in the design of theranostic tools. Having support for the hypothesis that the mimotope library peppos, sampling major sequence clusters, is optimal when compared to a set of 8 other libraries, we next studied its diagnostic potential using sera from a larger set of patients (n=34) with brain tumors. Due to the small data set, the main goal was a “proof of principle” test demonstrating the capacity of the assay to provide mimotope profiles (feature subsets in machine learning parlance) suitable for building predictors for randomly selected pathology. The distribution of patients by diagnosis (glioblastoma multiforme – GBM, lung cancer metastases in the brain – ML, breast cancer metastases in the brain – MB, and non-tumor bearing patients – C) is shown in Table 3. After cleaning, local, global normalization and balancing the group sizes which warranted the use of ComBat [37] for the following batch compensation, the reactivity data represented 28 patients’ serum IgM binding to 586 peptides. The comparison between the staining intensities of the mimotopes (or features) in the patients’ diagnostic groups yielded overlapping sets of reactivities significantly expressed in each diagnosis as compared to the other two - 290 features for GBM, 263 for ML, and 204 for C. Overall, 380 features showed significant reactivity in at least one of the diagnostic groups. The “negative” peptides (library pepnegrnd) represented 49/206 non-significant and 24/380 significant reactivities (χ^2^, p<0.0001). The finding of individuals with IgM reactive for some of them when testing a larger group is not a surprise. That is why the background reactivity was considered more reliably determined by the data analysis, rather than on the mean level of the pepnegrnd library.

**Table 3.**
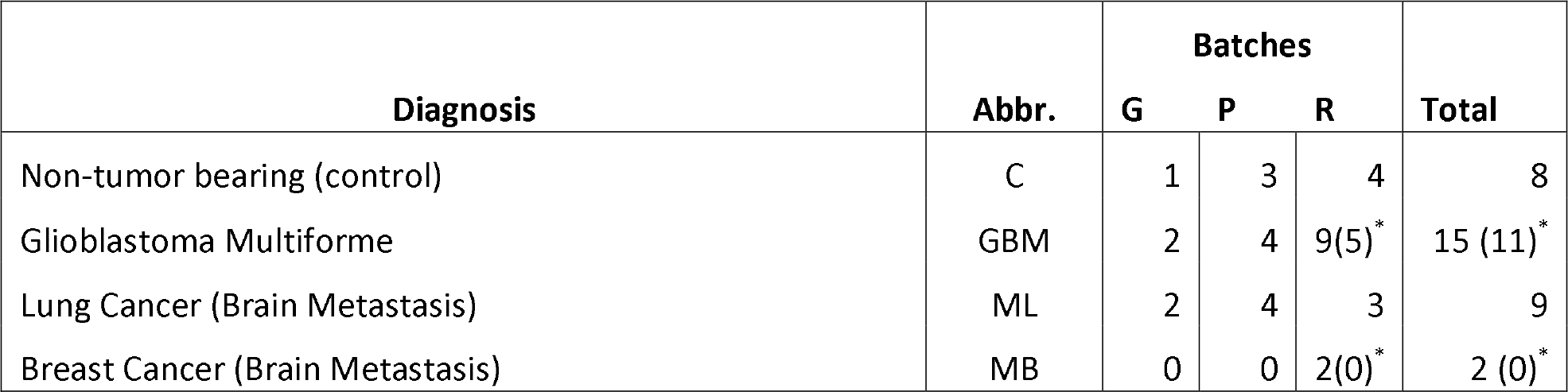
Patients tested using the optimized library.

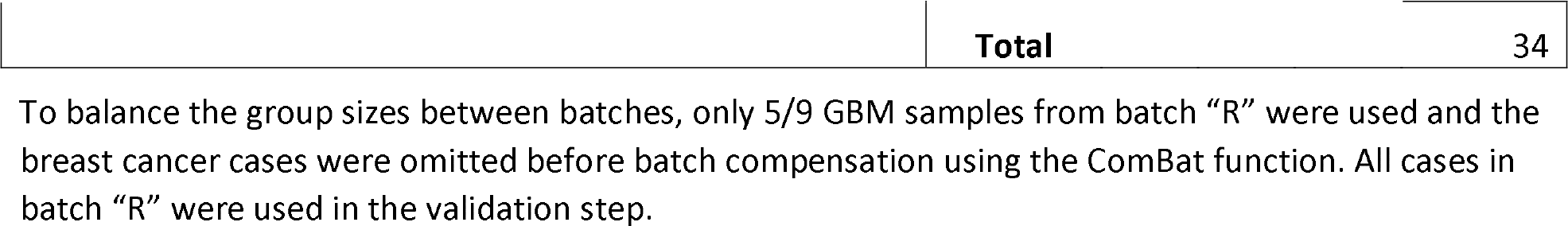

A projection of the cases on the 380 positive reactivities by multidimensional scaling (MDS) which maps the data to two dimensions showed no separation (Suppl. Fig.9). The feature space is highly multidimensional. The peptide library is not targeted to any particular pathology, but represents a universal tool for IgM repertoire studies. Therefore, a feature selection step is necessary to construct a predictor for each diagnostic task.

A recursive elimination algorithm was applied whereby features were removed successively in a way that improves the separation of the patient data clusters of interest until no further improvement of separation is possible (see Supplemental methods section for details). Using this approach, we tested the capacity of the smaller feature sets, thus selected, to separate dichotomously GBM from the rest. Support vector machine (SVM) models based on these optimal feature subsets still suffered from overfitting as demonstrated by a leave one out validation (data not shown). Aiming at a better generalization, next we explored the variation of the feature sets selected by recursive elimination using sets of patient data that differed by two cases in a bootstrap scheme (Supplemental Methods). It was surprising to find that so similar teaching sets differed considerably in the optimal features (mimotope reactivities) selected by them with only 4 features common for all patient sets. The reason for this could be the variability between individuals and the capacity of the mimotope library to reflect it. It was possible to demonstrate that the best prediction both of the teaching and of the validation sets was achieved when using the features that recurred in at least 50% of the bootstrap runs of the recursive elimination algorithm (Fig. 7).

**Figure 7.**
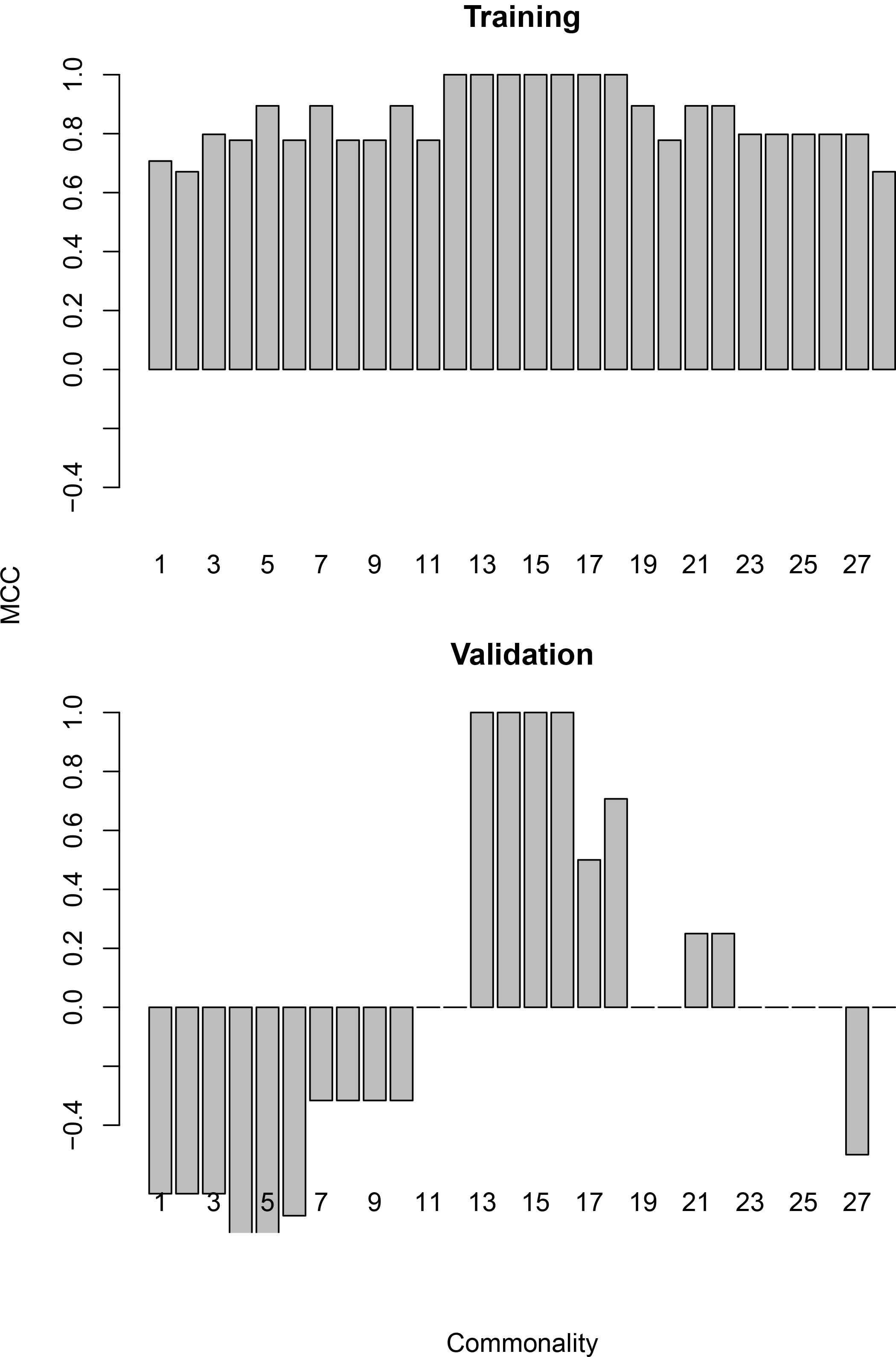
Matthew’s correlation coefficient as a measure of the prediction quality for SVM models constructed using GBM predicting feature sets of different minimal commonality. Minimal commonality of n means that the features in the set are found in n or more of the bootstrap sets. The validation set consists of the cases in batch “R” that were omitted from the batch compensated united sets. The model predicts these cases as belonging to the same class as the rest of the respective cases in batch “R”. Since the values in batch “R” were not subject to batch compensation the validation also serves as a control against confounding introduced by the ComBat function.

Interestingly, this two stage feature selection strategy helped improve considerably the generalization and a SVM model constructed on a 2-dimensional mapping (using multidimensional scaling) of the IgM reactivity to the set of 55 mimotopes, thus selected, successfully classified the GBM and non-GBM cases in the validation set of sera.

**Figure 8.**
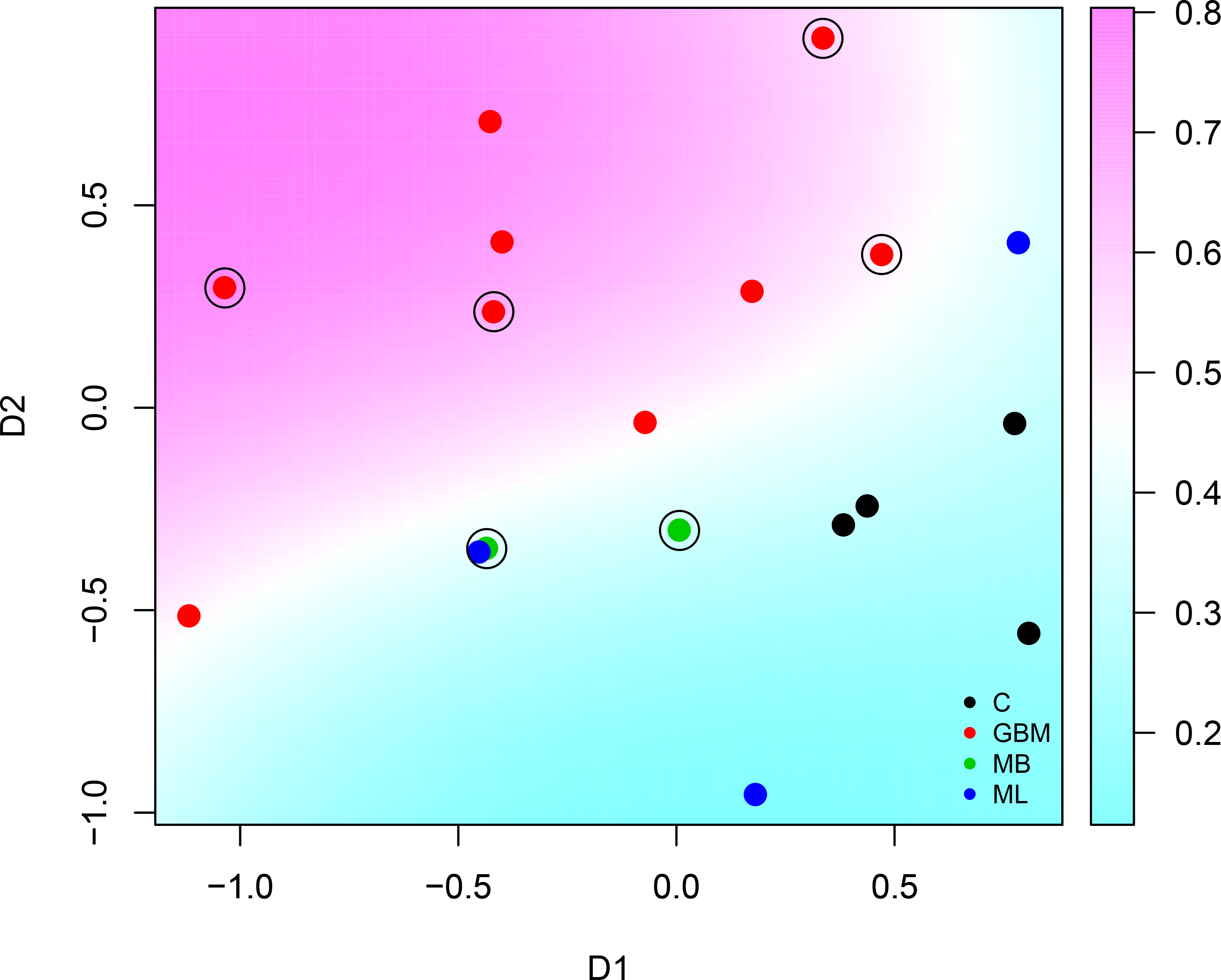
Multidimensional scaling plot of cases in batch “R” based on the feature set of minimal commonality of 50%. See figure 7 and Supplemental Methods for details. The encircled points correspond to the validation set.

Thus, we were able to show that a rationally designed small library of 586 IgM mimotopes contains potentially a huge number of mimotope profiles that can differentiate randomly selected diagnoses after appropriate feature selection.

## Discussion

High-throughput omics screening methods have led to the identification of biomarkers as profiles extracted from a particular dynamic diversity – proteome, genome, glycome, secretome, etc. The use of the antibody repertoire as a source of biomarkers has been defined and approached in multiple ways. First came the technically minimalistic, but conceptually loaded, semi quantitative immunoblotting, developed 20 years ago. This technique served as no less than a paradigm setter for systems immunology [38–43]. The further development produced methods that have been referred to as functional immunomics [33] in terms of protein reactivities, as immunosignaturing [31] in terms of random peptide libraries, or described as a deep panning technique [44] and in terms of igome of mimotopes selected from random phage display libraries. Here we describe the design of the first mimotope library for the analysis of the human IgM repertoire of reactivities recurrent in most individuals [12,45,46].

The deep panning approach relies on next generation sequencing (NGS), and thus requires balancing between sequence fidelity and diversity. Even with diversity affected by discarding sequences of one and two copies on the one hand, and overgrowth of phage clones on the other, our strategy still manages to find a general representation of the mimotope sequence space by identifying clusters of mimotopes. This relatively small set of sequence classes is hypothesized to be related to the modular organization of the repertoire defined previously [47].

The central role of prolines in the natural antibody mimotopes has been observed previously [48]. Tchernychev et al. also used a phage display library, and now it is clear that the high proline content is related to the bias of the particular phage display library. This property of the library may facilitate the discovery of mimotopes because prolines are associated with turns and flanking structures and proline abundance also reduces the entropic component of the binding. The selection by the IgM repertoire led to an enrichment of tryptophan and negatively charged residues in the middle of the sequences suggesting that the public IgM reactivity has a preference for loop like mimotopes (facilitated by the presence of prolines) with negative charges. The abundance of tryptophan is also interesting in terms of its propensity (together with proline) to mimic carbohydrate structures [49].

The mimotope library of diversity 10^5^ derived by deep panning reflects the recurrent (also referred to as public) IgM specificities found in the human population. The low IgM reactivity to the library of random peptides with sequences least related to the 790 clusters suggests that not only the library of a little over 200 000 mimotopes represents well the IgM mimotope space but also that the 790 sequence clusters provide a rather complete description of that space. The good coverage of the IgM reactivity space may be facilitated by the polyspecific binding of IgM to small peptides.

Although the large mimotope library can be used as is in large arrays when applicable it is not very practical for routine diagnostics. The classification in 790 clusters was used to produce a smaller and more applicable library for clinical use, of a subset of approx. 600 mimotopes (peppos) by picking representative sequences from the most significant clusters. Thus, this library was designed and shown to optimally represent the mimotopes’ main public reactivity patterns found in the phage selection experiment. The proposed optimized small library could be used as a tool for the study of the IgM public repertoire, as a source of mimotopes for design of immunotherapeutics [50–53], but mostly it may by applied as a multipurpose diagnostic tool.

As a diagnostic tool, the optimized small library has some key properties that distinguish it from other omic sets. Since it is designed to represent practically ubiquitous public specificities, the sets of features (mimotope reactivities), significantly expressed in the different diagnoses, were overlapping considerably. No single reactivity was correlating strongly with a whole diagnostic group, but subsets of reactivities collectively could separate the diagnoses. Thus, feature selection becomes essential for the design of predictors based on polyspecificities. Using the proposed algorithms, the typical feature set tuned for a dichotomous separation of diagnoses contained between 28 and 111 sequences (median=66). The improvement of generalization by keeping only features recurring in the bootstrap feature selection algorithm helped reduce the overfitting of the models. The optimal feature set for GBM diagnosis contained 55 mimotopes. Thus, if the library provides in the order of 500 significant reactivities, the theoretical capacity of this approach is >10^70^ different subsets in case of just qualitative differences of presence or absence of reactivity. Thus, the information provided by a typical IgM binding assay with the library is probably enough to describe any physiological or pathological state of clinical relevance reflected in the IgM repertoire. Of course, this is just an estimate of the resolution of the method. The number of naturally occurring profiles and their correlation with clinically relevant states will determine the actual capacity.

The novelty of our approach is based on the combination of several previously existing concepts:

First, early studies argued that the physiologically autoreactive natural antibodies comprise a consistent, organized immunological compartment [40,43,54–57]. The consistency of the natural antibody self-reactivity among individuals was considered evidence for the existence of a relatively small set of preferred self-antigens. Such “public reactivities” are most probably related to the germline repertoire of antibodies generated by evolutionarily encoded paratope features and negative/positive selection [34,58]. These antibodies were targeted using protein microarrays, the utility of which has been previously demonstrated [23,33,34,47]. Recently, the existence of structurally distinct public V-regions has been analyzed using repertoire sequencing [12], noting that they are often found in natural antibodies. If the repertoire should be read as a source of information providing consistent patterns that can be mapped to physiological and pathological states, the public natural IgM autoreactivity seems to be a suitable but underused compartment.

Second, germline variable regions are characterized by polyspecificity or cross-reactivity with protein and non-protein antigens [14]. It seems that going for epitopes could be a way to approach the repertoire convolution. Yet, the actual epitopes will be mostly conformational and hard to study. In similar tasks, mimotopes are often used [59–62], yet M.H. Van Regenmortel argues that mimotopes are of little use to structural prediction of the B-cell epitope [61]. Therefore, their utility might be rather in the structural study of the repertoire as a whole.

Third, the usage of peptide arrays for the analysis of the antibody repertoire is increasingly popular [44,63–66]. It involves the use of random peptide arrays for extracting repertoire immunosignatures by some groups and deep panning of phage display libraries to analyze antibody responses by others. Since an antibody can often cross-react with a linear epitope that is part of the nominal conformational epitope [61], the 7-residue library offers suitable short mimotopes as compared to typical B-cell epitope combining exhaustive set of sequences with considerable structural complexity. Furthermore, from the Immunoepitope Database (http://www.iedb.org) collection of linear B cell epitopes, 4821 of 45829 entries are less than 8 residues long.

The library also provides a rich source of mimotopes that can be screened for different theranostic tasks focused on particular targets. On an omics scale, the smaller optimized mimotope library proposed here probes efficiently the relevant repertoire of public IgM reactivities matching its dynamic diversity with potentially over 10^70^ distinct profiles. The major task ahead is designing studies aimed at efficiently extracting specific diagnostic profiles and building appropriate predictors, e.g. – for classifying immunotherapy responders, predicting the risk of malignancy in chronic inflammation, etc.

## Materials and methods

### Deep panning

Human IgM was isolated from a sample of IgM enriched IVIg - IgM-Konzentrat (Biotest AG, Dreieich, Germany, generously provided by Prof. Srini Kaveri), while human monoclonal IgM paraprotein was isolated from an IgM myeloma patient’s serum selected from the biobank at the Center of Excellence for Translational Research in Hematology at the National Hematology Hospital, Sofia (with the kind cooperation of Dr. Lidiya Gurcheva). In both cases, IgM was purified using affinity chromatography with polyclonal anti-μ antibody coupled to agarose (A9935, SIGMA-ALDRICH, USA). A 7-mer random peptide library (E8100S, Ph.D. -7, New England Biolabs, USA) was panned overnight at 4°C on pooled human IgM adsorbed on polystyrene plates at a concentration of 0.1 mg/ml, washed, eluted with glycine buffer at pH 2.7 and immediately brought to pH7. The eluate was transferred to a plate coated with monoclonal IgM and incubated according to the same protocol, but this time the phage solution was collected after adsorption and amplified once, according to Matochko et al. [35]. Briefly, the phage DNA was extracted and the peptide-coding fragment amplified by PCR. The amplicons were subjected to deep sequencing using the Next Seq platform (Illumina, USA), performed at the Sequencing Core Facility of Oslo University Hospital.

### Patients’ sera

Sera from randomly selected patients with glioblastoma multiforme (GBM), low grade glioma (G), brain metastases of breast (MB) or lung (ML) cancers, as well as non-tumor bearing patients (C) (herniated disc surgery, trauma, etc.) of the Neurosurgery Clinic of St. Ivan Rilski University Hospital, Sofia acquired according to the rules of the ethics committee of the Medical University in Sofia, after its approval and obtaining informed consent, were analyzed on the sets of peptides defined in microarray format. The sera were aliquoted and stored at −20°C. Before staining the sera were thawed, incubated for 30 min at 37°C for dissolution of IgM complexes, diluted 1:100 with PBS, pH 7.4, 0.05% Tween 20 with 0.1% BSA, further incubated for 30 min at 37°C and filtered through 0.22μm filters before application on the chips.

### Peptide microarray

The customized microarray chips were produced by PEPperPRINT™ (Heidelberg, Germany) by synthesis in situ as 7-mer peptides attached to the surface through their C-terminus and a common spacer GGGS. The layout was in a format of a single field of up to 5500 or five fields of up to 600 peptides in randomly positioned duplicates. The chips were blocked for 60 minutes using PBS, pH 7.4, 0.05% Tween 20 with 1% BSA on a rocker, washed 3×1 min with PBS, pH 7.4, 0.05% Tween 20 and incubated with sera in dilutions equivalent to 0.01 mg/ml IgM (approx. 1:100 serum dilution) on a rocker overnight at 4°C. After 3×1 minute washing the chips were incubated with secondary antibodies at RT, washed, rinsed with distilled water and dried by spinning in a vertical position in empty 50 ml test tubes at 100 × g for 2 minutes.

### Microarray data treatment

The microarray images were acquired using a GenePix 4000 Microarray Scanner (Molecular Devices, USA). The densitometry was done using the GenePix^®^ Pro v6.0 software. All further analysis was performed using publicly available packages of the R statistical environment for Windows (v3.4.1) (Bioconductor – Biostrings, limma, pepStat, sva, e1071, Rtsne, clvalid, entropy, RankProd, multcomp) as well as in house developed R scripts (https://github.com/ansts/IgMimoPap1 and https://github.com/ansts/IgMimoPap2). For algorithm details see Suppl. Methods.

## Supporting information

Sequence logos of the t-sne clusters of underrepresented sequences

High definition version of Fig. 6

Position weighted matrices of 790 clusters of mimotopes

## Competing interests

The authors declare no competing interests.

## ACKNOWLEDGEMENTS

This work was performed with the support of EEA/Norway Grant BG09/D03-103 and the Bulgarian Fund for Scientific Research Grant D01-11/2016. The authors wish to thank Prof. Radha Nagarajan, Prof. Ivanka Tsakovska and Prof. Soren Hairabedyan for critically reading the manuscript and a number of useful comments.

## Author Contribution

A.P. conceptualized the project, analyzed the results performing all the in silico work, supervised experiments except for the sequencing as well as the overall project execution and prepared the manuscript;

M. Hadzhieva ran the phage display experiments;

V.K. and M. T. ran the microarray experiments up to data processing, catalogued and maintained the seroteque;

V.S. supervised the phage display experiments, participated in the conceptualizing the paper and together with M. Heinz and L.A.M.Z. carried out the DNA isolation, PCR and sequencing;

E.H. supervised the sequencing task, participated in conceptualizing the project and the preparation of the manuscript;

S. P. and M.T. performed the data processing of microarray scans;

T.V. and T.K.E participated in conceptualizing the project, analysis of the results and the preparation of the manuscript;

D.F. was responsible for the patient selection, informed consent, ethics committee protocol preparation, blood collection and serum preparation.

