## Supplementary figures and images for "A rationally designed mimotope library for profiling of the human IgM repertoire"

### High definition version of Fig. 6

Mixture of Selected Mimotopes and Random Peptides

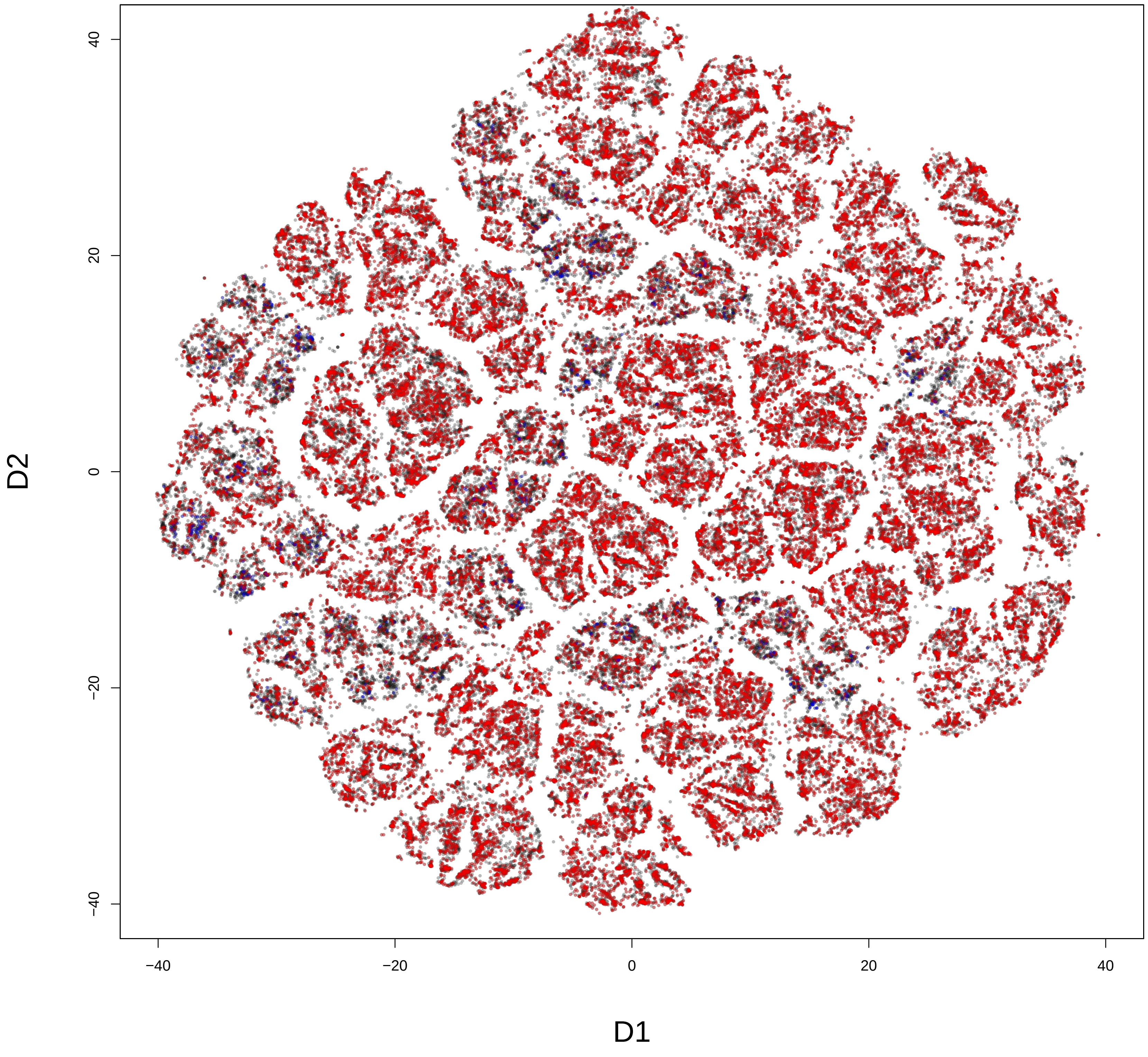


Fig. 8

### Sequence logos of the t-sne clusters of underrepresented sequences

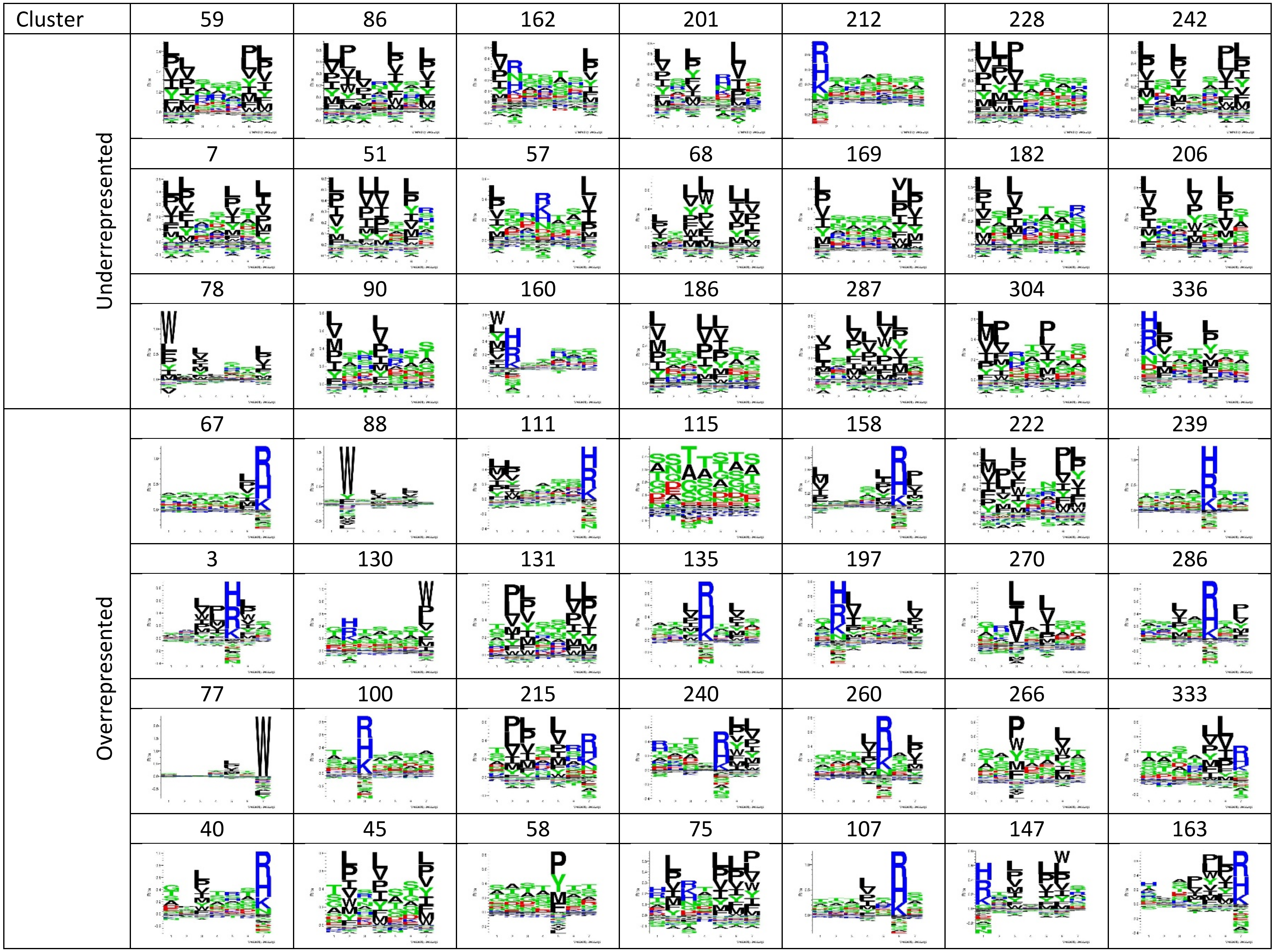
