## Supplemental tables for "A rationally designed mimotope library for profiling of the human IgM repertoire"

Supplement Tables

**Suppl. Table 1.** GLM results from comparison between mean peptide reactivity in different libraries (for library designation see Table 1).

Simultaneous Tests for General Linear Hypotheses

Multiple Comparisons of Means: Tukey Contrasts

| Linear Hypotheses: | Estimate | Std. Error | z value | Pr(>\|z\|) | |
|---|---|---|---|---|---|
| pep5pred - pep5 == 0 | 0.040685 | 0.036587 | 1.112 | 0.9502 | |
| pepneg - pep5 == 0 | -0.101131 | 0.039385 | -2.568 | 0.1563 | |
| pepneglo - pep5 == 0 | -0.001002 | 0.075444 | -0.013 | 1 | |
| pepnegrnd - pep5 == 0 | -0.390134 | 0.036239 | -10.766 | < 0.0010 | *** |
| pepother5 - pep5 == 0 | 0.059812 | 0.03348 | 1.786 | 0.6110 | |
| peppos - pep5 == 0 | 0.081682 | 0.03848 | 2.123 | 0.3810 | |
| peprnd - pep5 == 0 | -0.115647 | 0.036324 | -3.184 | 0.0286 | * |
| pepneg - pep5pred == 0 | -0.141816 | 0.037469 | -3.785 | 0.0035 | ** |
| pepneglo - pep5pred == 0 | -0.041687 | 0.074462 | -0.56 | 0.9992 | |
| pepnegrnd - pep5pred == 0 | -0.430819 | 0.034148 | -12.616 | < 0.0010 | *** |
| pepother5 - pep5pred == 0 | 0.019127 | 0.031204 | 0.613 | 0.9985 | |
| peppos - pep5pred == 0 | 0.040997 | 0.036517 | 1.123 | 0.9477 | |
| peprnd - pep5pred == 0 | -0.156332 | 0.034237 | -4.566 | < 0.0010 | *** |
| pepneglo - pepneg == 0 | 0.100129 | 0.075876 | 1.32 | 0.8832 | |
| pepnegrnd - pepneg == 0 | -0.289003 | 0.037129 | -7.784 | < 0.0010 | *** |
| pepother5 - pepneg == 0 | 0.160943 | 0.034442 | 4.673 | < 0.0010 | *** |
| peppos - pepneg == 0 | 0.182813 | 0.03932 | 4.649 | < 0.0010 | *** |
| peprnd - pepneg == 0 | -0.014516 | 0.037212 | -0.39 | 1 | |
| pepnegrnd - pepneglo == 0 | -0.389132 | 0.074292 | -5.238 | < 0.0010 | *** |
| pepother5 - pepneglo == 0 | 0.060814 | 0.072986 | 0.833 | 0.9903 | |
| peppos - pepneglo == 0 | 0.082684 | 0.07541 | 1.096 | 0.9539 | |
| peprnd - pepneglo == 0 | -0.114645 | 0.074333 | -1.542 | 0.7706 | |
| pepother5 - pepnegrnd == 0 | 0.449946 | 0.030795 | 14.611 | < 0.0010 | *** |
| peppos - pepnegrnd == 0 | 0.471816 | 0.036168 | 13.045 | < 0.0010 | *** |
| peprnd - pepnegrnd == 0 | 0.274487 | 0.033865 | 8.105 | < 0.0010 | *** |
| peppos - pepother5 == 0 | 0.02187 | 0.033404 | 0.655 | 0.9978 | |
| peprnd - pepother5 == 0 | -0.175459 | 0.030895 | -5.679 | < 0.0010 | *** |
| peprnd - peppos == 0 | -0.197329 | 0.036253 | -5.443 | < 0.0010 | *** |

---
Signif. codes:  0 ‘***’ 0.001 ‘**’ 0.01 ‘*’ 0.05 ‘.’ 0.1 ‘ ’ 1
(Adjusted p values reported -- single-step method)

**Suppl. Table 2.** GLM results from comparison of total correlation between peptide reactivity in different libraries (for library designation see Table 1).

```
Simultaneous Tests for General Linear Hypotheses

Multiple Comparisons of Means: Tukey Contrasts


Fit: glm(formula = log(value) ~ Var2, data = mc10clKLD)

Linear Hypotheses:
                            Estimate Std. Error  z value Pr(>|z|)    
pep5pred - pep5 == 0        0.055920   0.001796   31.138   <0.001 ***
pepneg - pep5 == 0          0.048193   0.001796   26.835   <0.001 ***
pepnegrnd - pep5 == 0       0.058865   0.001796   32.778   <0.001 ***
pepother5 - pep5 == 0      -0.032593   0.001796  -18.148   <0.001 ***
peppos - pep5 == 0         -0.316416   0.001796 -176.188   <0.001 ***
peprnd - pep5 == 0          0.033380   0.001796   18.587   <0.001 ***
pepneg - pep5pred == 0     -0.007728   0.001796   -4.303   <0.001 ***
pepnegrnd - pep5pred == 0   0.002945   0.001796    1.640    0.656    
pepother5 - pep5pred == 0  -0.088513   0.001796  -49.286   <0.001 ***
peppos - pep5pred == 0     -0.372337   0.001796 -207.326   <0.001 ***
peprnd - pep5pred == 0     -0.022540   0.001796  -12.551   <0.001 ***
pepnegrnd - pepneg == 0     0.010673   0.001796    5.943   <0.001 ***
pepother5 - pepneg == 0    -0.080785   0.001796  -44.983   <0.001 ***
peppos - pepneg == 0       -0.364609   0.001796 -203.023   <0.001 ***
peprnd - pepneg == 0       -0.014812   0.001796   -8.248   <0.001 ***
pepother5 - pepnegrnd == 0 -0.091458   0.001796  -50.926   <0.001 ***
peppos - pepnegrnd == 0    -0.375282   0.001796 -208.966   <0.001 ***
peprnd - pepnegrnd == 0    -0.025485   0.001796  -14.191   <0.001 ***
peppos - pepother5 == 0    -0.283824   0.001796 -158.040   <0.001 ***
peprnd - pepother5 == 0     0.065973   0.001796   36.735   <0.001 ***
peprnd - peppos == 0        0.349797   0.001796  194.775   <0.001 ***
---
```

**Suppl. Table 3.** GLM results from comparison between mean correlation between patient profiles in different libraries (for library designation see Table 1).

```
Simultaneous Tests for General Linear Hypotheses

Multiple Comparisons of Means: Tukey Contrasts


Fit: glm(formula = value ~ variable, data = corptzm)

Linear Hypotheses:
                            Estimate Std. Error z value Pr(>|z|)
pep5pred - pep5 == 0        0.010342   0.027400   0.377  0.99995
pepneg - pep5 == 0         -0.024782   0.027400  -0.904  0.98574
pepneglo - pep5 == 0       -0.114978   0.027400  -4.196  < 0.001 ***
pepnegrnd - pep5 == 0      -0.059773   0.027400  -2.181  0.36269
pepother5 - pep5 == 0      -0.003980   0.027400  -0.145  1.00000
peppos - pep5 == 0         -0.133697   0.027400  -4.879  < 0.001 ***
peprnd - pep5 == 0         -0.051258   0.027400  -1.871  0.57126
pepneg - pep5pred == 0     -0.035123   0.027400  -1.282  0.90583
pepneglo - pep5pred == 0   -0.125319   0.027400  -4.574  < 0.001 ***
pepnegrnd - pep5pred == 0  -0.070115   0.027400  -2.559  0.17162
pepother5 - pep5pred == 0  -0.014322   0.027400  -0.523  0.99955
peppos - pep5pred == 0     -0.144039   0.027400  -5.257  < 0.001 ***
peprnd - pep5pred == 0     -0.061600   0.027400  -2.248  0.32313
pepneglo - pepneg == 0     -0.090196   0.027400  -3.292  0.02225 *
pepnegrnd - pepneg == 0    -0.034991   0.027400  -1.277  0.90753
pepother5 - pepneg == 0     0.020801   0.027400   0.759  0.99505
peppos - pepneg == 0       -0.108915   0.027400  -3.975  0.00182 **
peprnd - pepneg == 0       -0.026476   0.027400  -0.966  0.97906
pepnegrnd - pepneglo == 0   0.055204   0.027400   2.015  0.47168
pepother5 - pepneglo == 0   0.110997   0.027400   4.051  0.00127 **
peppos - pepneglo == 0     -0.018719   0.027400  -0.683  0.99744
peprnd - pepneglo == 0      0.063720   0.027400   2.325  0.28004
pepother5 - pepnegrnd == 0  0.055793   0.027400   2.036  0.45724
peppos - pepnegrnd == 0    -0.073924   0.027400  -2.698  0.12324
peprnd - pepnegrnd == 0     0.008515   0.027400   0.311  0.99999
peppos - pepother5 == 0    -0.129717   0.027400  -4.734  < 0.001 ***
peprnd - pepother5 == 0    -0.047278   0.027400  -1.725  0.67064
peprnd - peppos == 0        0.082439   0.027400   3.009  0.05339 .
---
Signif. codes:  0 ‘***’ 0.001 ‘**’ 0.01 ‘*’ 0.05 ‘.’ 0.1 ‘ ’ 1
(Adjusted p values reported -- single-step method)
```

**Suppl. Table 4.** GLM results from comparison of mean nearest neighbor distance between pepetide profiles in different libraries (for library designation see Table 1).

```
Simultaneous Tests for General Linear Hypotheses

Multiple Comparisons of Means: Tukey Contrasts


Fit: glm(formula = log(value) ~ L1, data = mc10clnndist)

Linear Hypotheses:
                             Estimate Std. Error z value Pr(>|z|)
pep5pred - pep5 == 0       -0.0463163  0.0204399  -2.266  0.29596
pepneg - pep5 == 0         -0.0314013  0.0220028  -1.427  0.83374
pepneglo - pep5 == 0        0.2372490  0.0421481   5.629  < 0.001 ***
pepnegrnd - pep5 == 0      -0.0576799  0.0202455  -2.849  0.07630 .
pepother5 - pep5 == 0      -0.0583229  0.0187041  -3.118  0.03474 *
peppos - pep5 == 0          0.0708564  0.0214974   3.296  0.01993 *
peprnd - pep5 == 0         -0.0260619  0.0202927  -1.284  0.89734
pepneg - pep5pred == 0      0.0149150  0.0209328   0.713  0.99629
pepneglo - pep5pred == 0    0.2835652  0.0415995   6.817  < 0.001 ***
pepnegrnd - pep5pred == 0  -0.0113636  0.0190771  -0.596  0.99882
pepother5 - pep5pred == 0  -0.0120067  0.0174327  -0.689  0.99701
peppos - pep5pred == 0      0.1171727  0.0204009   5.744  < 0.001 ***
peprnd - pep5pred == 0      0.0202544  0.0191272   1.059  0.96175
pepneglo - pepneg == 0      0.2686503  0.0423893   6.338  < 0.001 ***
pepnegrnd - pepneg == 0    -0.0262786  0.0207429  -1.267  0.90381
pepother5 - pepneg == 0    -0.0269216  0.0192415  -1.399  0.84765
peppos - pepneg == 0        0.1022577  0.0219666   4.655  < 0.001 ***
peprnd - pepneg == 0        0.0053394  0.0207890   0.257  1.00000
pepnegrnd - pepneglo == 0  -0.2949288  0.0415043  -7.106  < 0.001 ***
pepother5 - pepneglo == 0  -0.2955719  0.0407746  -7.249  < 0.001 ***
peppos - pepneglo == 0     -0.1663925  0.0421291  -3.950  0.00185 **
peprnd - pepneglo == 0     -0.2633108  0.0415273  -6.341  < 0.001 ***
pepother5 - pepnegrnd == 0 -0.0006431  0.0172043  -0.037  1.00000
peppos - pepnegrnd == 0     0.1285363  0.0202060   6.361  < 0.001 ***
peprnd - pepnegrnd == 0     0.0316180  0.0189192   1.671  0.68953
peppos - pepother5 == 0     0.1291794  0.0186614   6.922  < 0.001 ***
peprnd - pepother5 == 0     0.0322611  0.0172599   1.869  0.55321
peprnd - peppos == 0       -0.0969183  0.0202533  -4.785  < 0.001 ***
---
Signif. codes:  0 ‘***’ 0.001 ‘**’ 0.01 ‘*’ 0.05 ‘.’ 0.1 ‘ ’ 1
(Adjusted p values reported -- single-step method)
```
