## Supplemental figures for "A rationally designed mimotope library for profiling of the human IgM repertoire"

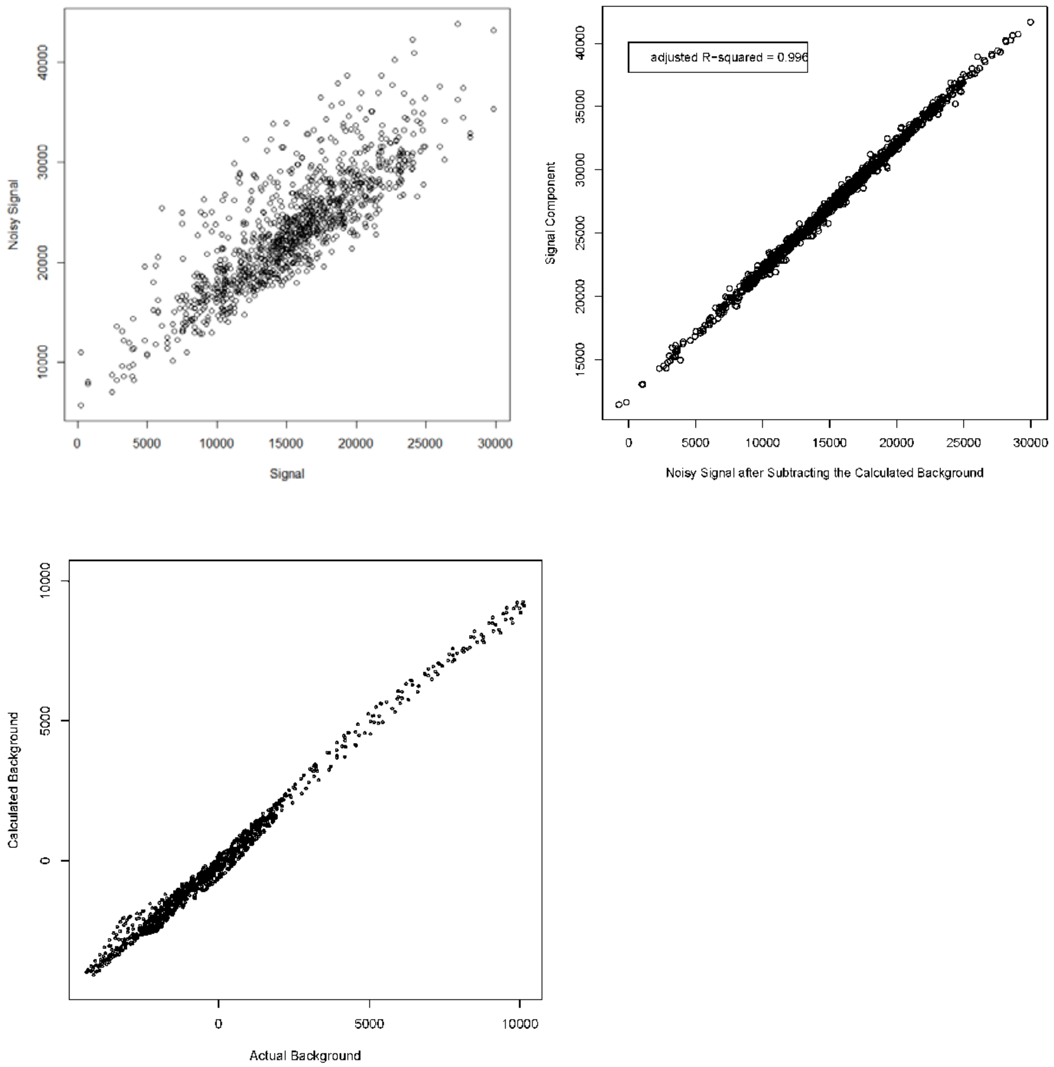


**Suppl. Figure 1**. Results of a simulated example of background subtraction for microarray data local normalization using the randomly positioned spot duplicates.

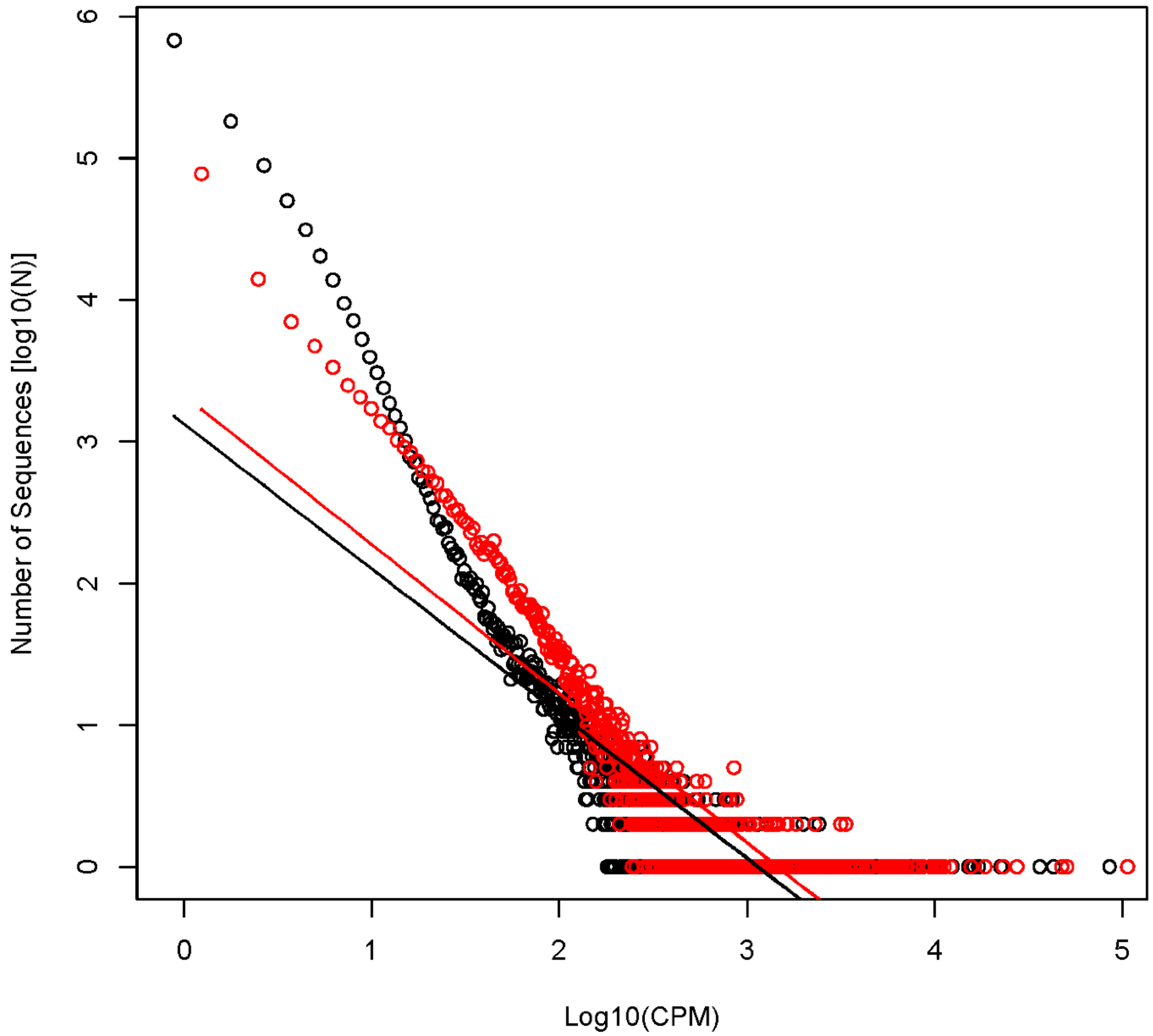


**Suppl. Figure 2**. Results from NGS. Plot of reads distribution by CPM (counts per million). Black – sample 1 – selected from original library, Red – sample 2 – selected form a pre-amplified original library. Here the selection caused deviation from the power law (the straight lines) with an emphasis on the lower CPM. The pre-amplification enriches in better fit phages and leads to a considerable shrinkage of the highly diverse compartment of low CPM where most of the targeted diversity lies.

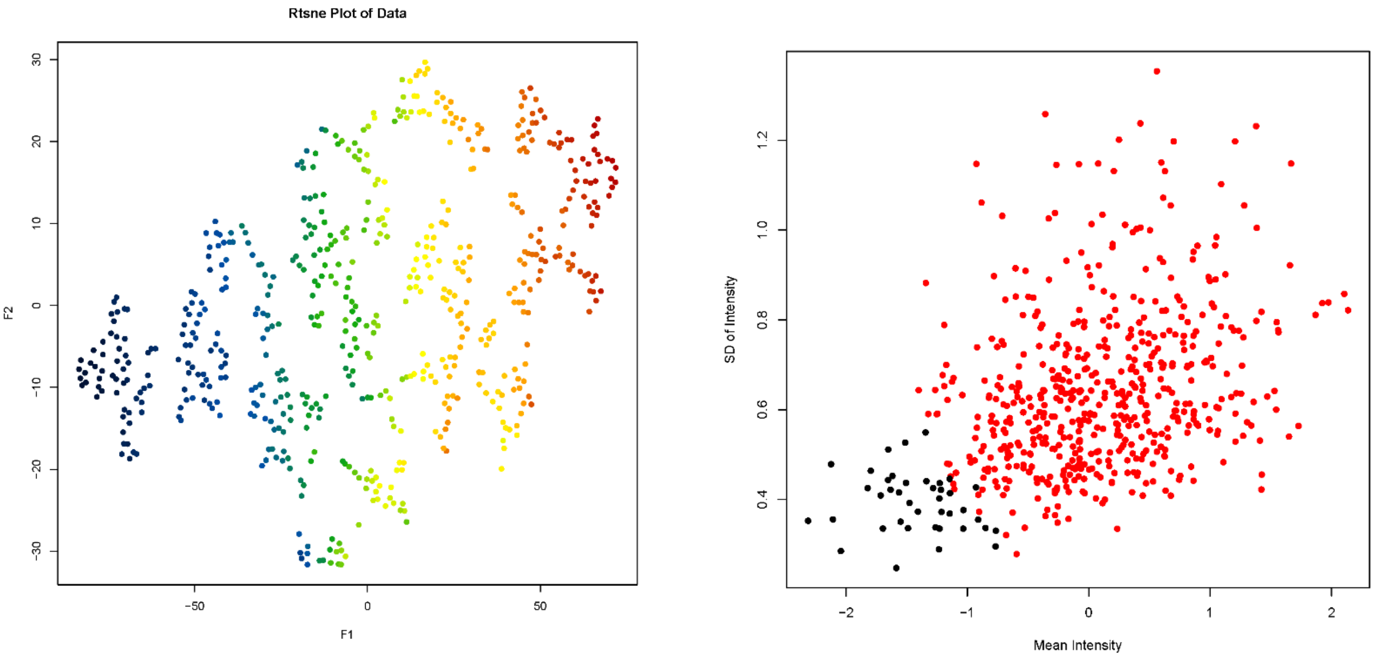


**Suppl. Figure 3.** Defining background binding by clustering using t-sne. The clustering separated a group of low binding peptides (left panel - color coded for mean binding). Right panel – the actual SD by mean data color coded for the sequences belonging to the same cluster.

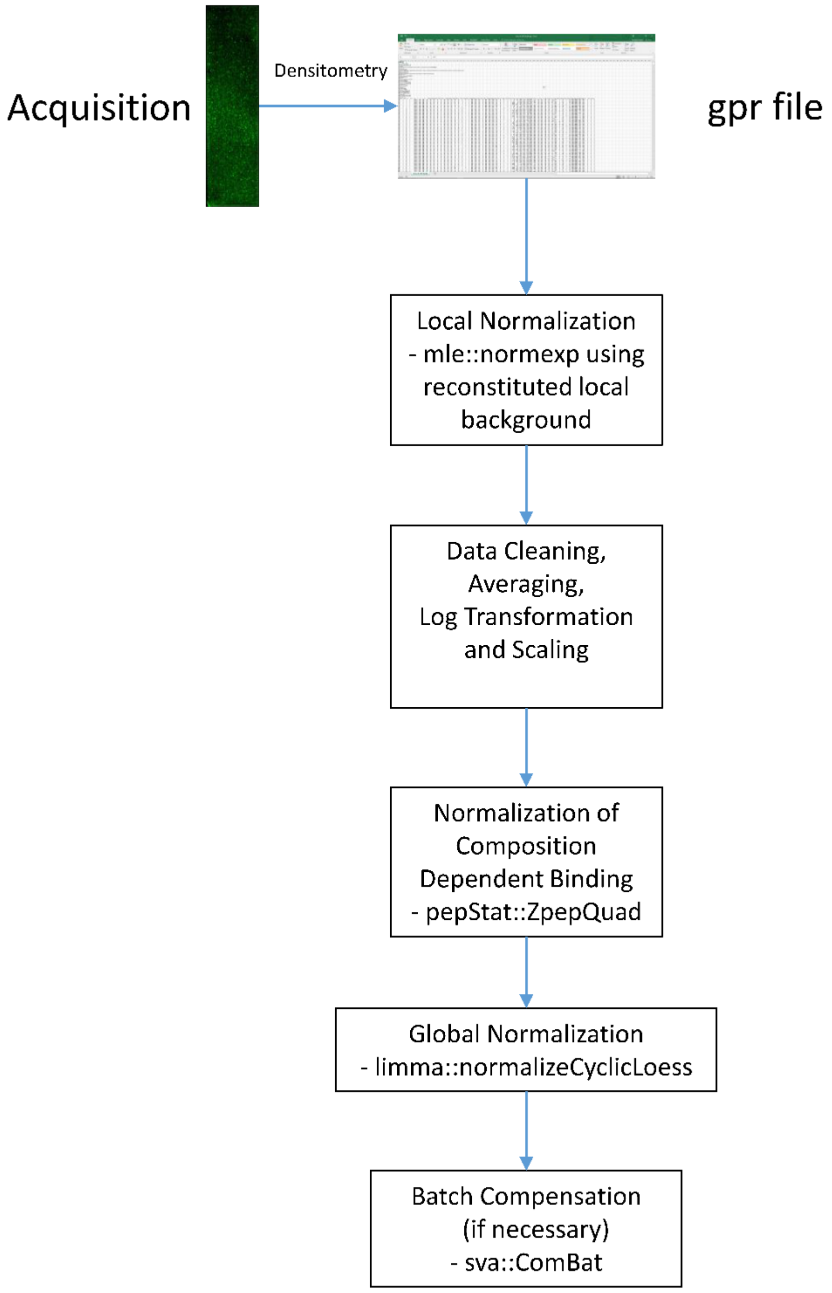


**Suppl.Figure 4.** Workflow for the microarray analysis.

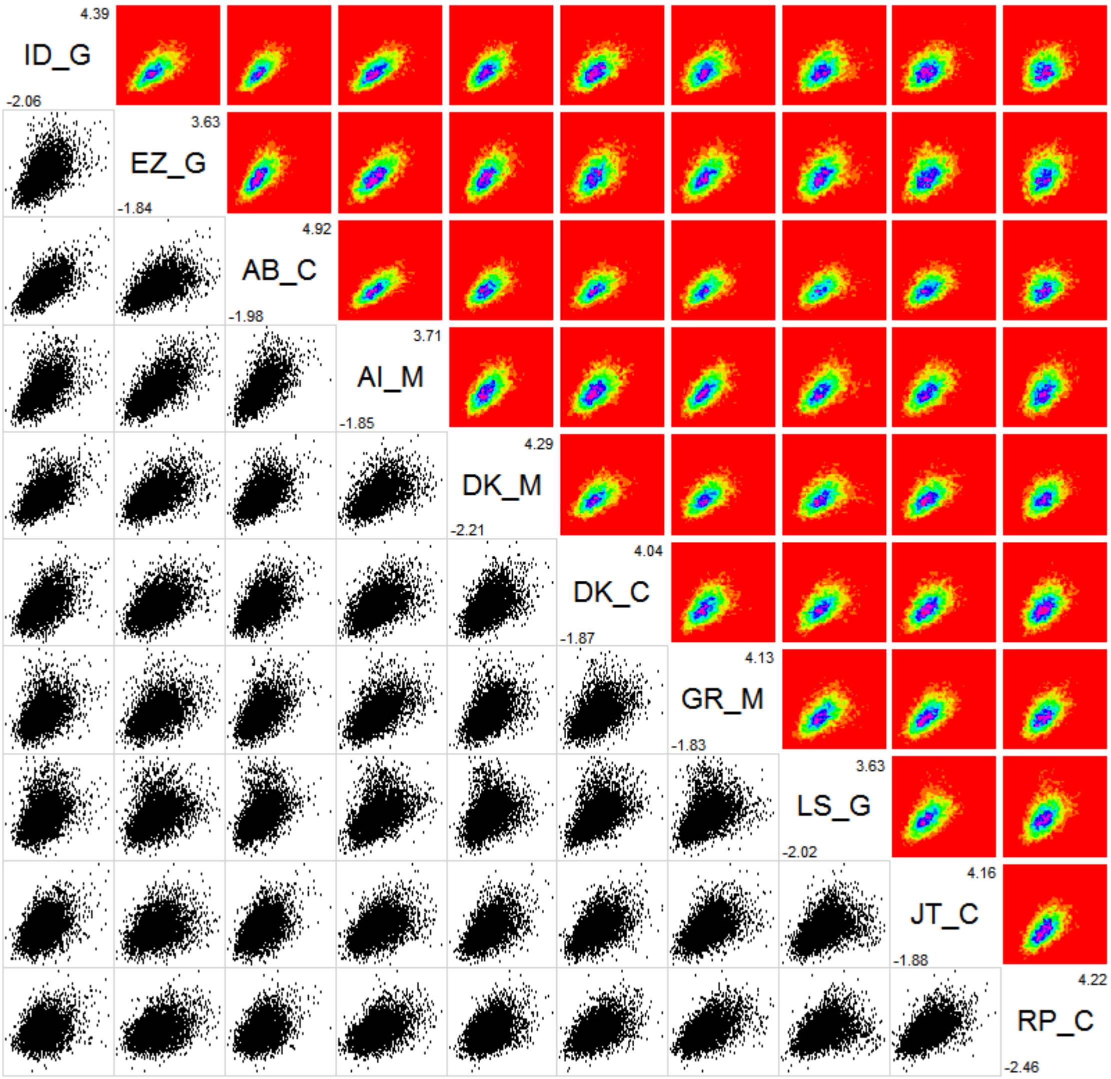


**Suppl. Figure 5.** Correlogram of data from the binding of patient serum IgM to a collection of 5500 peptides representing all the studied libraries. The data is log transformed and scaled and is plotted after local and global normalization.

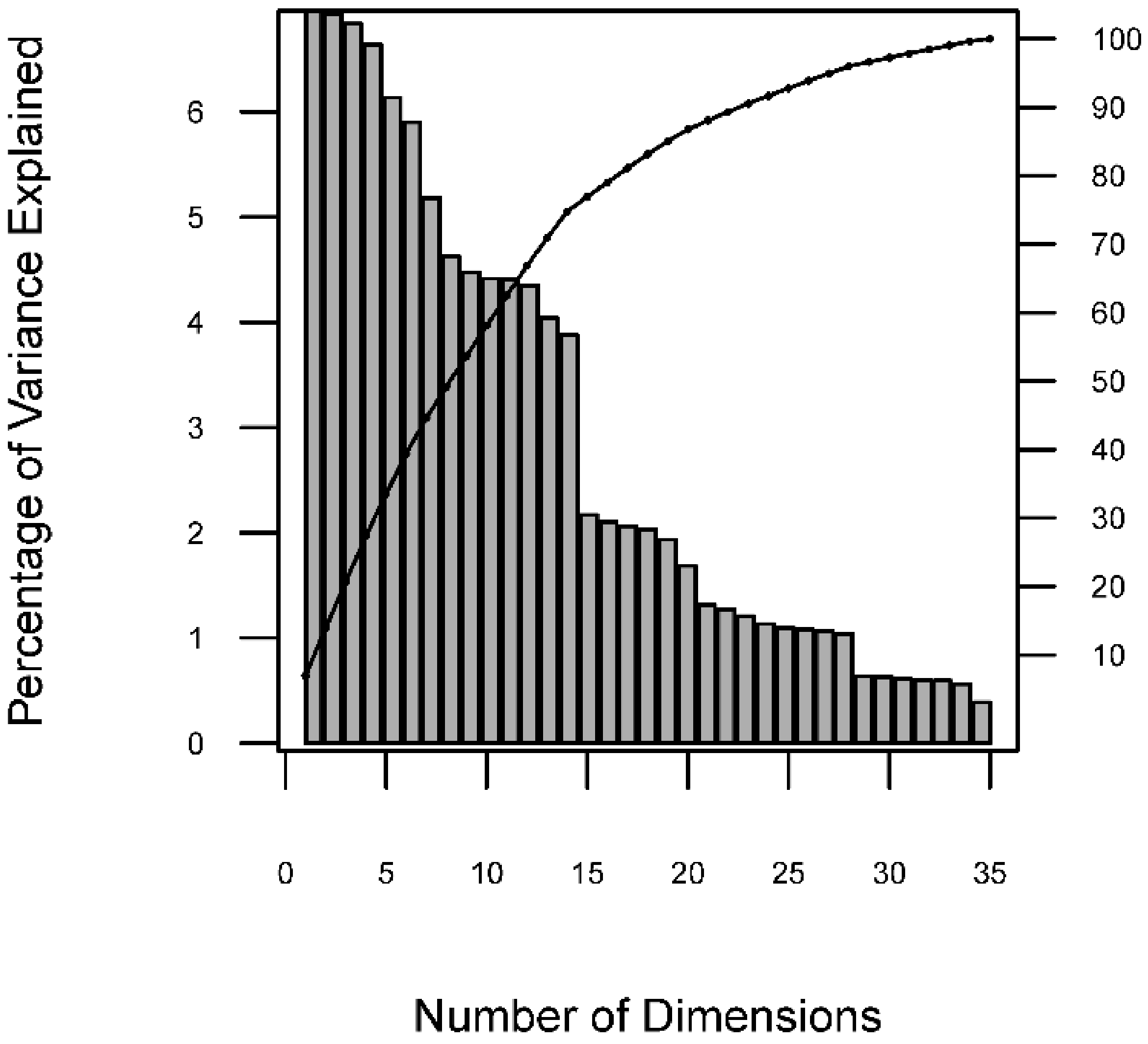


**Suppl. Figure 6.** Scree plot for the PCA on the phage selected mimotope library encoded using the five dimensional z-scores as described in Materials and Methods. The cumulative variance plot is associated with the right axis.

A

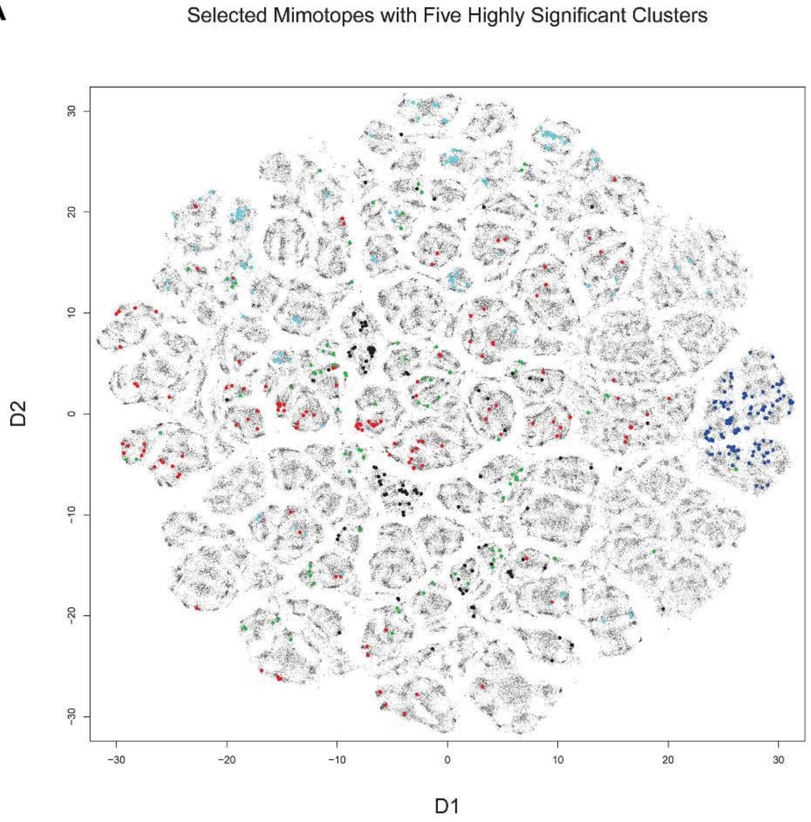

Selected Mimotopes with Five Highly Significant Clusters
D2
D1
30
20
10
0
-10
-20
-30


B

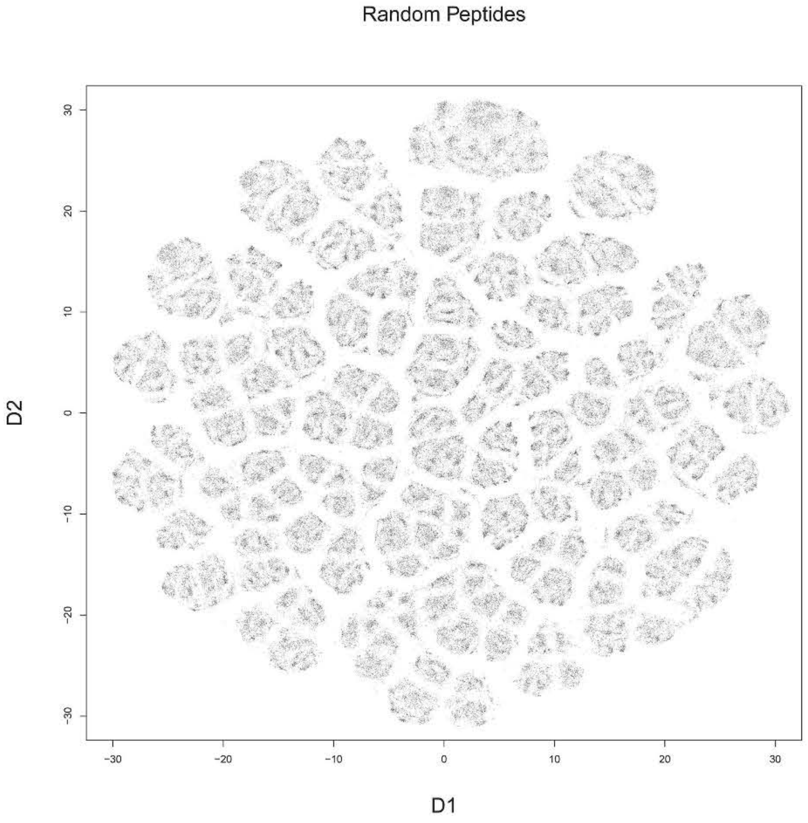

Random Peptides
D2
D1
30
20
10
0
-10
-20
-30

**Suppl. Figure 7.** Visualization of the 7-mer sequence space of the phage display selected mimotopes (A) and equal number random sequence with the same background residue frequencies (B) constructed using t-sne based on the Barnes-Hut algorithm. The peptides in five clusters (included in library pep5) are color coded in (A). The peptide sequences were encoded using a 5-dimensional score reflecting basic biophysical properties of the amino acids. Each sequence is represented by a 35 dimensional vector but the t-sne mapping is performed after initial reduction to 14 dimensions by PCA. There is only moderate correlation between the clustering visualized by t-sne and the GibbsCluster classification. The image is of high resolution and can be zoomed for better detail inspection.

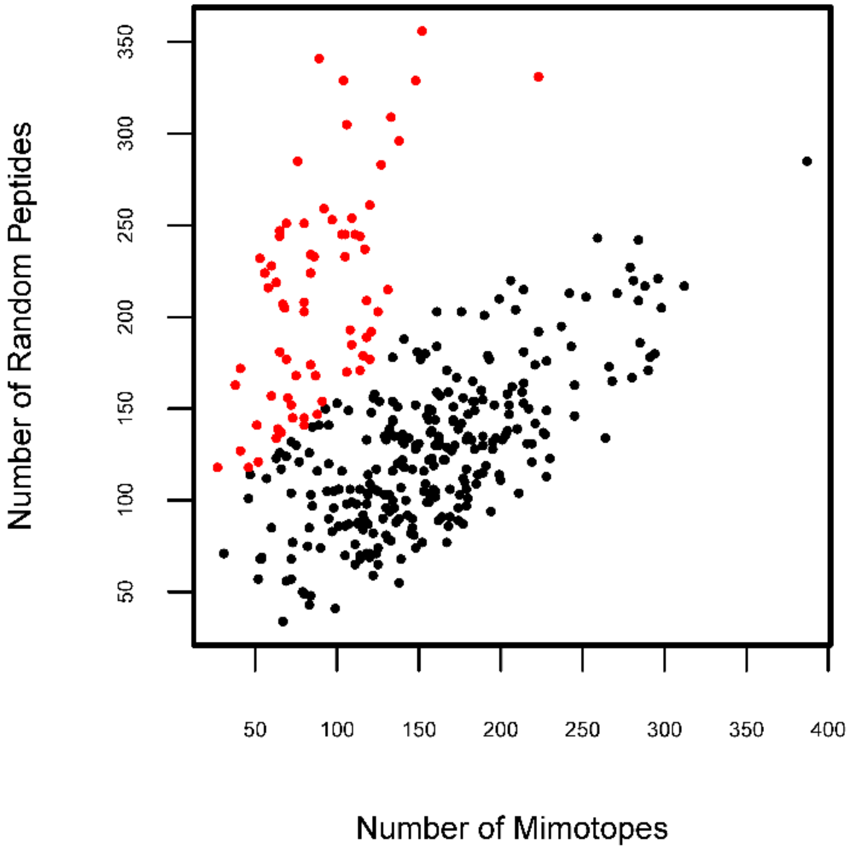


**Suppl. Figure 8.** Number of random or mimotope peptides per cluster in the t-sne map. Each point represents one cluster characterized by the number of mimotopes in it (X) or – the number of random sequences (Y). A mixture of 2 two dimensional normal distributions was fitted and the points corresponding to clusters under-represented by the mimotopes are colored red (probability to belong to the alternative component of the distribution p<0.1).

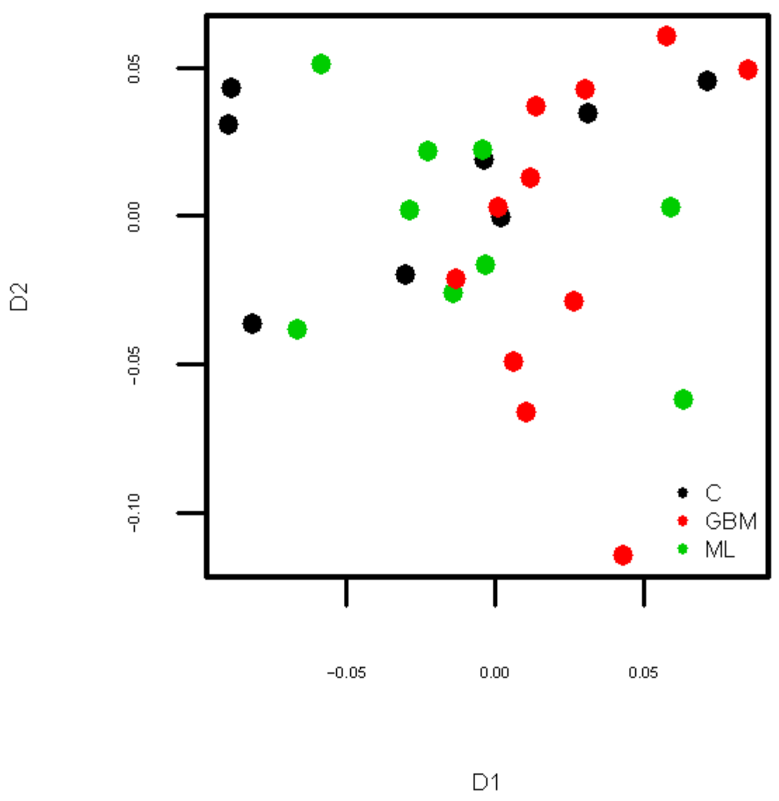


**Suppl. Figure 9.** Multidimensional scaling plot of the patient data projected on the 380 features significantly expressed in any of the diagnostic groups. No separation is observed due probably to the diversity of the individual repertoires reflected in multitude of profiles of mimotope reactivities. More specific diagnostic sets should be isolated by feature selection.

**Suppl. Figure 10.** Plots of the clustering criterion as a function of the number of selected features generated by the recursive elimination feature selection algorithm. The search starts with all features and successively eliminates the feature that results in the greatest increase of the clustering criterion (the process proceeds from the far end back to less features). After an optimal set of features is found (at the maximum of the curve) the further elimination of even the most unnecessary feature leads to a fall in the criterion value. Thus, the feature set corresponding to the maximum of the curve is taken as the optimal one. Each curve corresponds to the dichotomous clustering of one diagnosis against all other cases: black – controls, red – GBM, blue – ML, green – MB. Three random curves out of 28 are shown for each of the diagnoses.

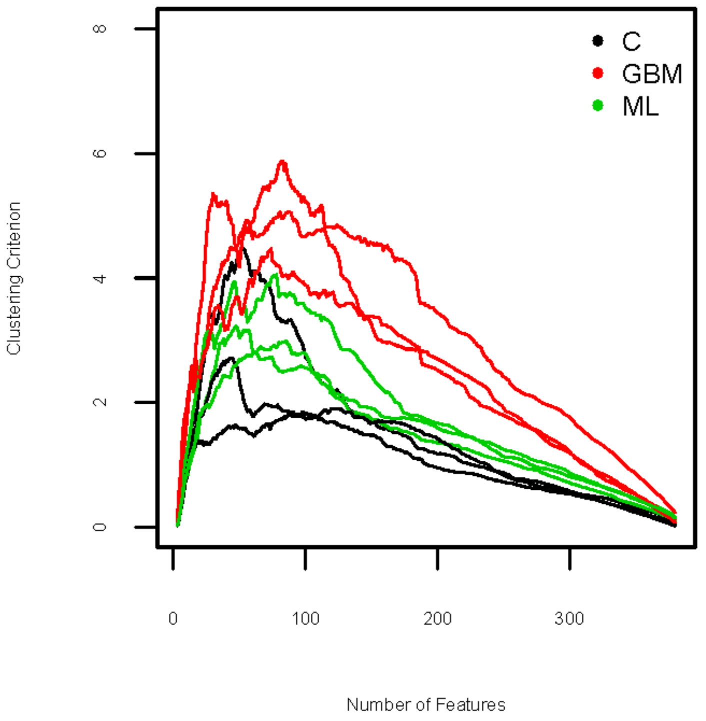

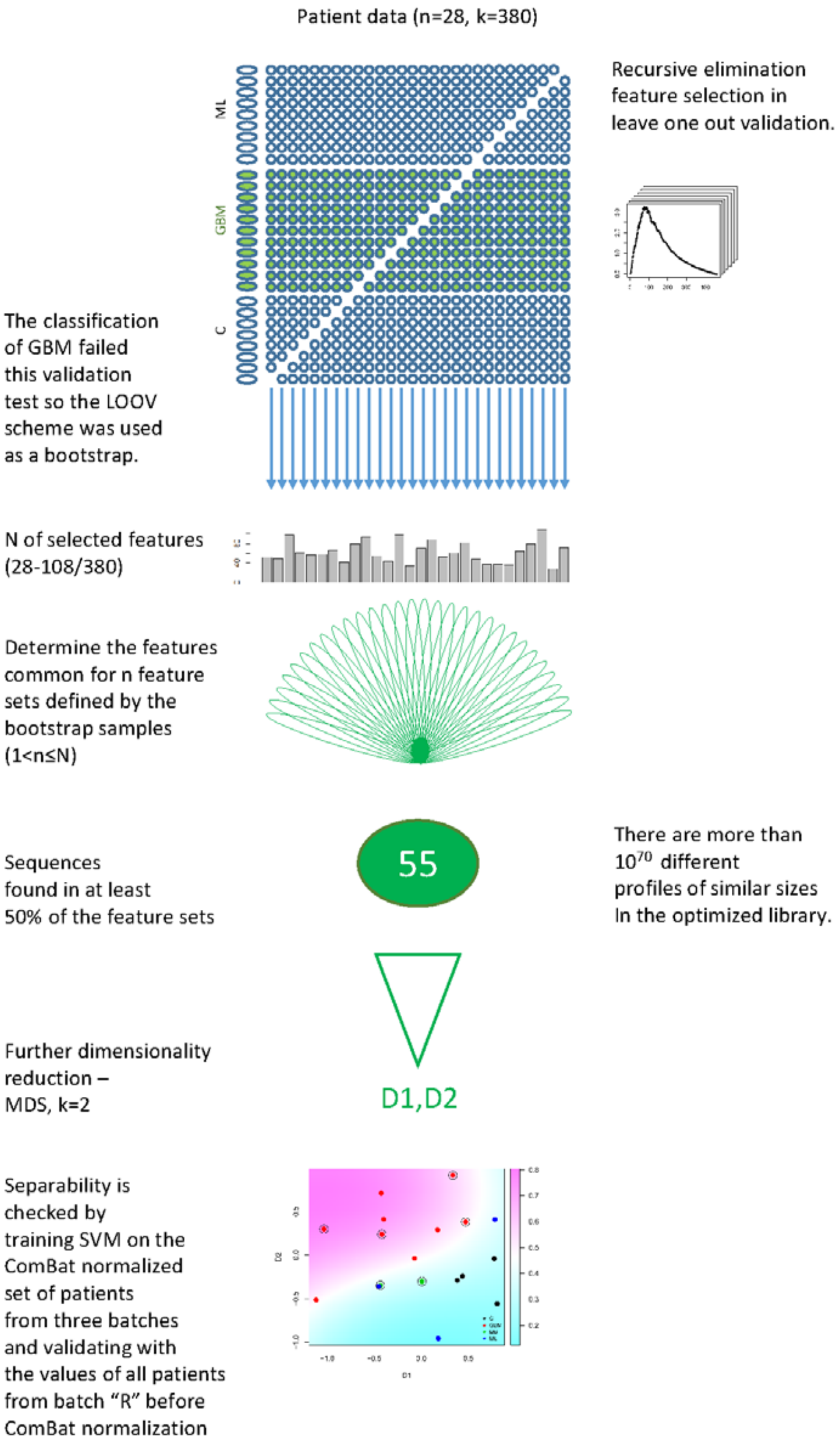


**Suppl. Figure 11.** Schematic representation of the feature selection scheme for extracting diagnostic profiles and a SVM model.
