## Supplemental methods for "A rationally designed mimotope library for profiling of the human IgM repertoire"

**Microarray local normalization**

For the initial analysis comparing different libraries the approximation was based on a set of spots comprising the lower tertile of intensities determined locally in overlapping patches of 5x5 spots. For the rest of the chips the background approximation was done using a new method based on the randomly positioned duplicates.

The method is based on the fact that by definition the duplicate spots should have the same signal intensity $I$ and different local background levels $B$ because they are positioned randomly over the whole area of the chip. Thus, the difference between the intensities $\Delta I$ of the duplicate spots of a given peptide is independent of the peptide's signal intensity $S$ but equals the gradient of the background $\Delta B$ between the two duplicate distant spots plus an error term $e$:

$$I = S + B + e \quad (1)$$

$$\Delta I = \Delta B + e \quad (2)$$

Let's consider a spot $I_{i0,j0}$. The spots $I_{i,j}$ ($i=i0\text{-}2..i0+2$, $j=j0\text{-}2..j0+2$) adjacent to $I_{i0,j0}$ have all very similar background levels unlike their respective duplicates which generally lie away. The average of the differences between each spot's intensity in the patch and its respective duplicate $\overline{\Delta I_{i0,j0}} = \frac{\sum \Delta \boldsymbol{I_{i,j}}}{\mathbf{9}}$ approximate the background at $I_{i0,j0}$ centered to its mean. Indeed, for a random variable $X$ with values $X_i$ ($i=1..n$) the mean of the differences $X_i$-$X_j$ ($j=1:n$) equals the centered $X_i$ value. When $j$ is only a sample rather than all the values the mean differences are still an approximation of the centered $X_i$ value with an error increasing with the decreasing size of the sample. To improve the approximation, the calculated background values were iteratively adjusted by $(\overline{\Delta I_{i0,j0}} - \Delta I_{i0,j0})/s$ where $\Delta I_{i0,j0}$ is the observed duplicate difference at $I_{i0,j0}$ and s is a step factor empirically adjusted to 2. The approximated centered background correlated with the real in a simulated example (Suppl. Fig. 1).

with $R^2$=0.99. Using overlapping patches has also the effect of smoothing the calculated background. The iteration typically converges after about 20 cycles.

**Mimotope selection**

The reads from the deep sequencing experiment were processed using the script provided by Matochko et al. [1]. The Phred quality score cut off used was 32 with a probability for an erroneous base call of $6x10^{-4}$. The unique reads from experiments A and B were pooled so that only one copy of a sequence existed in the final set (n=1 100 124). These were further filtered based on the number of occurrences of each sequence. The criterion for retaining a sequence was an occurrence in 3-10 CPM, yielding 224 087 unique sequences using the following rationale. Considering that 78% of the single base changes lead to a change in the encoded amino acid [2] (see also phD7GC script for the mean substitution rate in Ph.D-7 library), the used Phred score led to a frequency of 0.00328 for the sequences with one erroneous amino acid residue if there is only one wrong base call per sequence. The occurrence of two or more errors in the same read is in the order of $10^{-5}$ and will be considered negligible. Since the criteria for inclusion is at least 3 CPM, the probability for all 3 to be erroneous is $3.5x10^{-8}$. If at least one of the reads is an actually existing mimotope the presence of wrong calls in the triplet is non-consequential. Thus, the error rate of the mimotope calls is negligible at Phred score of 32.

The limit of 10 copies on the high copy numbers was applied after comparison of the distributions of the number of clones by CPM between the original and the preamplified library (Suppl. Fig. 2). It showed that the threshold of 10 copies was discriminating the original and preamplified libraries, with diversity in the latter skewed towards highly proliferating clones. This fact was interpreted in view of the observation that the affinity selection seemed to favor low CPM clones. Therefore, the high CPM clones were excluded to avoid a possible contamination with non-selected clones having an advantage when they are highly proliferating. This restriction led to the exclusion of 9.96% of the reads. Reads found in 3-10 copies were selected yielding 224087 sequences which contained none of the parasitic sequences reported by Matochko et al. [3]. This mimotope library was further subjected to clustering using the GibbsCluster-2.0 method [4]. The number of clusters was optimized using the Kulback-Leibler distance (KLD) from the background model of random sequences [4]. Position weighted matrices (PWM) were defined for each cluster using pseudo counts as follows:

$$PWM_{k,j} = log_2\left(\frac{\sum I\left(X_{i,j} = k\right) + b_k\sqrt{N}}{(N + \sqrt{N})b_k}\right)$$

where i=1..N are the rows of the alignment, j=1:7 are the columns of the alignment, I(aa=k) is the indicator function used to count the occurrences of amino acid k in column j and $b_k$ is the background frequency of amino acid k in the phage display library [5].

Using the PWMs, the median of the log odds (LO) scores of the peptides in each cluster was calculated. Next, the probability of the occurrence of peptides with LO greater than the respective median in a set of random peptides 10-fold larger than the analyzed library was determined empirically. Using this estimate, the probability of the chance occurrence of as many peptides with scores higher than the median score as observed in each cluster was calculated using the binomial distribution. The probabilities, thus found for each cluster, were used as a criterion for their relevance to the mimotope library being generated.

For filtering random peptide sequences for sequences related to the mimotope library, each of $2.3x10^6$ random 7-mer peptide sequences was tested against each of the PWM of mimotope clusters defined. For each random peptide, only the score of the top scoring cluster was retained, and the peptides were ranked in the ascending order of these scores. The lowest ranking peptides represented random sequences that were the least related to any of the clusters in the selected library were used.

The clustering of the mimotopes was visualized using the t-sne (t-Distributed Stochastic Neighbor Embedding) algorithm [6] in its implementation using the faster Barnes-Hut algorithm (Rtsne package) with theta parameter default value of 0.5 and a maximum of 1500 iterations [7].

The amino acid residues were described using the 5-dimensional scale of amino acid residue properties published by Hellberg et al. [8]. The amino acid residue properties quantification used was the same as for the pepStat binding normalization [9]. The 5 scales (z1-z5) were extracted as the latent variables describing the major factors underlying the variability of amino acids in the space of 26 physicochemical parameters. Thus, each peptide was represented by a vector of 35 scores corresponding to its seven positions. For a comparison, the same number of random 7-mer peptides generated using the amino acid background frequencies of Ph.D.-7 were clustered using the same parameters. The clusters in the t-sne plot were labeled using k-mean clustering.

**Microarray data treatment**

*Local normalization*

The spot intensities of a non-treated chip were subtracted from the stained chip spot intensities. Treatment with secondary antibody only did not result in any binding. Next, the local background had to be inferred due to the dense spot layout. To that end an approximated background was smoothed by support vector regression and the result passed to the backgroundCorrect [10] function for local normalization using the normexp (mle) method. The initial background approximation is described above.

*Global normalization*

The peptides with missing values (flagged “bad”) in some of the patients were removed. Log transformed locally normalized microarray data were next normalized for amino acid composition dependent binding using the ZpepQuad method of pepStat package [11]. This step is considered indispensable because of the strong effect of amino acid composition on binding due mostly to electrostatic interactions. The amino acid residue properties were quantified using the 5 dimensional descriptor z1-z5 of Sandberg et al. (1998)

[9]. This was followed by global normalization using normalizeCyclicLoess (method affy) from the package limma [10] and subjected to batch effect compensation using the ComBat [12] function from package sva whenever necessary (for different chip batches and for different channels). The data was acquired in 2 different batches using the green (batch G) and the red channel (batch R). Eleven cases were part of an earlier experiment and that subset of the data could be used also in this assay (batch P). For batch effect compensation the groups were balanced by selecting a subset of the patients with relatively even representation of each tested diagnosis. The criterion for inclusion of the 5 GBM patients from batch "R" was the minimal difference of the mean and coefficient of variation from the mean and CV of the whole group of GBM in that batch. Finally, each peptide binding intensity was represented by the mean of its duplicates.

Because of a lack of a suitable negative control, the baseline binding was determined from the data. The clean data used for the testing of the diagnostic potential was subjected to dimensionality reduction of the peptide reactivities using t-sne [6], which clearly outlined a group of peptides with uniformly low reactivities, (Suppl. Fig. 3) that was considered background binding. The mean of the background binding intensity was subtracted from the data before testing for significant reactivities.

*Library comparison*

A general linear model, followed by Tukey contrast, was used to compare the expression of reactivities in the different tested libraries. The total correlation between the different peptide reactivity profiles of each library with 10 patients' serum IgM was used as a measure of the redundancy of the library. The profiles were viewed as points in a 10-mer space with each dimension corresponding to a patient. Total correlation was calculated as KLD from the theoretical maximal entropy joint distribution of the profile data, binned appropriately. To keep the number of bins comparable to the number of peptides, the intensity values for each patient were discretized just to "high" and "low" relative to the median for the studied library, effectively centering each library's data and, thus, removing the effect of the median intensity of each library. To further limit the number of bins, randomly selected bins (the same for all libraries) were aggregated in groups of 8, yielding 128 bins for the calculation of the baseline and the actual joint distributions. To equalize the size of the libraries, random samples of 380 peptides were used from each library, except for pepneglo which was excluded from this analysis due to its small size. The mean KLD values were determined in a bootstrap procedure with the described sampling repeated with replacement 100 times for the bin aggregation and 30 times for the library resampling producing 3000 samples. KLD for each library was thus calculated using the entropy::KL function.

The same approach could not be used for the transposed matrix, due to the disproportionately high number of bins necessary to describe the distribution in high/low values of 500-1000 different peptides. Instead, the mean correlation coefficient between the patient profiles across the peptides was used. To compare the mean correlation values between libraries by GLM, the correlation values were converted to z-scores.

Another test of the optimal sampling of the mimotope space was the mean nearest neighbor distance (NND) per library which was normalized (nNND) relative to the theoretical mean distance between the points in the 10-dimensional data cloud for the different libraries:

$$nNND_L = \frac{NND_L}{\left(\frac{V}{N_L}\right)^{\frac{1}{k}}},$$

where L=1..8, $N_L$ is the number of peptides in each library, k is the dimensionality (10 since the peptides are compared on the basis to their reactivity to 10 patients' sera) and V is the volume of the data cloud approximated as a k-dimensional ellipsoid [13]:

$$V = \frac{2\pi^{\frac{k}{2}}}{k\Gamma\left(\frac{k}{2}\right)} \prod_{i=1}^{k} (2 * \sigma_i),$$

where $\sigma_i$ is the standard deviation of the data along the $i_{th}$ dimension (the values of the $i_{th}$ serum) and $\Gamma$ is the gamma function. The logarithms of nNND were compared by general linear models (GLM).

Together with median intensity per library as a measure of the intensity of the signal, total correlation, mean correlation between patients' sera reactivities and mean nearest neighbor distance were combined to measure the optimization of the libraries. To make all four criteria positively correlated with the desired qualities of the libraries, the sign of the correlation measures was changed. The rank product method (p value for consistent "high expression" across the four tests) was used to test the optimization of the libraries.

**Visualization of the Mimotope Space**

T distributed stochastic neighbor embedding (t-sne) based on the Barnes-Hut algorithm was used to visualize the structure of the mimotope sequence space as represented by the general mimotope library produced by deep panning. The sequences were represented by converting each amino acid residue to a 5-dimensional vector of physical property scores as described in Materials and methods. Thus, each 7-mer peptide is represented by a 35-dimensional vector. The correlation dimension showed that the intrinsic dimension of the sequence space in this mapping was 11.25 for random peptides and 11.05 for the phage selected mimotope library. Principle component analysis indicated that the first 14 principle components account for approximately 75% of its variance (Supplemental Figure 6). The t-sne mapping was done after reducing the 35 dimensions to 14 by PCA (the initial.dims parameter of the Rtsne function).

To compare in more detail the mimotope space to the overall random sequence space, equally sized random samples of 50 000 mimotopes and random sequences were mapped together with the pepnegrnd library. After removing the duplicate sequences, the mixture contained 49964 mimotopes, 49950 random sequences and 684 peptides from the library pepnegrnd. This set of sequences was plotted again using the same approach as above (Suppl. Figure 7). The t-sne representation of the mixture of peptides was further clustered using k-means clustering in 350 clusters as a means of binning neighboring points. The number of random peptides and mimotopes were counted in each cluster (Supplement Figure 8).

**Feature selection algorithm**

The composition of the library underwent a small change - 75 of the lowest scoring sequences from pepnegrnd library were added to the library as a negative control. Because of the size limit of the final library, this addition was done by replacement of 75 of the selected positive peptides.

For the design of a practical algorithm for extracting information about a particular diagnosis, a feature selection approach was used based on recursive feature elimination. It used the quality of clustering of the predetermine diagnostic groups when mapped to only the selected subset of features as a criterion. The cluster separation of the cases of interest was measured using a combined clustering criterion:

$$Crit = \frac{100 * Dunn(d,c) * BH\gamma(d,c)}{Conn(d,c)}$$

Where Dunn(d,c) is the Dunn's clustering criterion [14,15] which is based on single inter-cluster and intra-cluster distances (extreme case), BHγ(d,c) is the Baker-Hubert Gamma index [16] which gives the overall agreement of distances and cluster assignment based on all the data, Conn is the connectivity validation measure for a given clustering partitioning based on 10 nearest neighbors , which emphasizes the agreement of distances and partitions in the vicinity of each point; *d* is the matrix of distances between the cases, and *c* is the diagnosis code for the cases. At each step the algorithm eliminates the feature the removal of which leads to the greatest increase in clustering criterion. This leads to an improvement of the clustering to a point at which the remaining features ensure optimal clustering. After that point the removal even of the feature that is least appropriate still leads to a decrease in the criterion. Supplemental Figure 10 shows the typical traces of the clustering criterion as a function of the number of features remaining with the best set of features corresponding to the maximum of the clustering criterion.

The recursive elimination features selection was performed on the union of features of significant expression in any diagnosis (n=380). The capacity of this approach to yield mimotope reactivity profiles that classify correctly the cases was tested in a leave one out validation scheme (LOOV) but it failed to generalize to the validation set probably due to the small training set.

In an attempt to improve the generalization of the prediction, next we used the data from LOOV as a bootstrap scheme testing the predictor's efficiency as a function of the number of bootstrap feature sets the features used appear in. The following algorithm was used for predicting GBM as contrasted to the rest of the cases. Let N be the number of cases. Each of the $N$ bootstrap samples contained the binding data for $N-1$ cases. The recursive elimination feature selection applied to each of those samples ultimately produced an intersecting family of $N$ feature sets. The common features were pooled by the number of sets they were common for (or "commonality") in groups with commonality greater than a threshold, i.e. – the group labeled 1 included all features, group n included features found in at least n bootstrap sets and group 28 contained the features found in all bootstrap sets. These groups were used to find the commonality level providing the best performing feature set.

The predictor was constructed after dimensionality reduction by MDS to two dimensions and using a radial basis function (Gaussian) kernel-based support vector machine. The performance of the model was measured using the Matthew's correlation coefficient (MCC). Among the set of features selected to differentiate the GBM cases, we used further those of 50% commonality (n=55) to predict the diagnoses of all the cases of batch "R". By including cases omitted before the batch compensation, this batch served both as a source of a validation set, as well as a control for the lack of confounding effect of the batch compensation. The Supplemental Figure 11 shows a schematic representation of the used feature selection algorithm.

**References to the Supplemental Methods**
